# The ABI5-WRKY45-LSU1 axis confers tolerance of *Arabidopsis thaliana* to cadmium

**DOI:** 10.64898/2026.04.20.718842

**Authors:** Fangjian Li, Xinni Zheng, Yanan Zhang, Jianjian Chen, Guihua Lv, Jinxiang Wang

**Author notes:** Correspondence: Corresponding Author Jinxiang Wang.

## Abstract

Abscisic acid (ABA) is involved in Cd tolerance in Arabidopsis, but the underlying mechanisms are unclear. In this study, we revealed that the ABI5-WRKY45-LSU1 axis confers the tolerance of Arabidopsis to Cd stress. Under Cd stress, the biosynthesis of ABA is increased, and the expression of transcription factor *ABI5* is upregulated. Accordingly, the *abi5-8* mutants show increased Cd sensitivity. ABI5 directly binds the ABRE element in the *WRKY45* promoter to activate its transcription. Overexpression of *WRKY45* rescues the Cd-hypersensitive phenotype of the *abi5-8* mutant, placing *WRKY45* downstream of *ABI5*. Transcriptome analyses identified *LSU1* as a potential *WRKY45* target. qRT-PCR, DUAL-LUC and EMSA experiments verified that WRKY45 binds the W-box *cis*-element in the *LSU1* promoter to activate its expression. Overexpression of *LSU1* enhances Cd tolerance by promoting the biosynthesis of non-protein thiols (NPT), glutathione (GSH), and phytochelatins (PC). Moreover, overexpression of *LSU1* suppresses Cd sensitivity in the *wrky45* mutant, confirming *LSU1* acts downstream of *WRKY45*. On the other hand, we found that ATP sulfurylase 1 (APS1) interacts with LSU1 based *in vitro* and *in vivo* evidences. LSU1 stabilizes APS1, slows its degradation, and enhances APS1 activity, thus leading to increased NPT, GSH, and PC accumulation and improved Cd detoxification. Notably, overexpressing *LSU1* did not rescue the Cd sensitivity of the *aps1-1* mutant, indicating that *LSU1* acts upstream of and depends on *APS1*. In short, we demonstrated a novel ABI5-WRKY45-LSU1 axis that regulates Cd tolerance through sulfur assimilation and phytochelatin synthesis.

**Highlights:**

1. Cadmium stress triggers ABA biosynthesis and *ABI5* expression; ABI5 directly binds to ABRE motifs in the WRKY45 promoter and activates its transcription.
2. WRKY45 transcriptionally activates LSU1, and LSU1 interacts with APS1 to stabilize it and elevate ATP sulfurylase activity, acting in an APS1-dependent manner.
3. The ABI5–WRKY45–LSU1 module enhances Arabidopsis Cd tolerance by boosting sulfur assimilation and GSH/PC-mediated Cd detoxification, rather than reducing Cd uptake.

## Introduction

Cadmium is heavy metal element that inhibit plant growth and development. Plants absorb Cd from soils and enter the human body through the food chain, endangering human health (Sahen et al.,2025). Cd toxicity can cause metabolic disorders in plant leaves, root browning, root length and dry weight reduction, further leading to cell death, and even plant death (Deng et al., 2021).

Plants have evolved elegant mechanisms to cope with Cd stress. These mechanisms include restricting Cd from entering cells, chelating Cd ions by anion groups, sequestrating the chelated Cd into vacuoles, and expelling Cd out of cells. The cell wall binds to Cd ions through negatively charged groups such as −OH, −COOH, etc., amino and aldehyde groups of pectin, reducing the entry of Cd ions into plant cells (Lwin et al., 2025). Glutathione (GSH) and phytochelatin (PC) are the main sulfur-containing ligands involved in Cd detoxification in plant cells (Wang et al., 2019). GSH itself or as a mercap to functional group in synthetic PC has a particularly strong affinity for Cd and can effectively adsorb Cd ions in the cytoplasm (Clemens, 2019). Vacuoles are the largest organelles in mature plant cells and play a key role in storing ions and metabolites, as well as detoxifying Cd. The HMA3 and ABCCs proteins transport Cd chelates, formed by GSH and PC, into the vacuole, reducing Cd toxicity (Hu et al., 2025).

Abscisic acid (ABA) is an important stress hormone that regulates physiological processes, antioxidant defense systems and the accumulation of osmotic regulators, helping plants cope with the adverse effects of Cd stress (Hu et al., 2020). As reported, exposure of plant roots to Cd-containing environments increases endogenous ABA levels in plant cells (Lu et al., 2020). Of note, being bZIP transcription factor, ABI5 is the crucial ABA signaling player. ABA upregulates *ABI5* expression (Li et al., 2025). ABI5 interacts with AtMYB49, inhibiting its function by preventing it from binding to the promoters of downstream genes *AtbHLH38*, *AtbHLH101*, *AtHIPP22* and *AtHIPP44*, resulting in reduced transcription levels of *AtIRT1* and reduced uptake of Cd (Lu et al., 2020). However, the functions of *ABI5* on Cd detoxification remain elusive.

The WRKY transcription factor is named after the WRKY domain, which contains approximately 60 amino acid residues and contains the conserved WRKYGQK sequence and a zinc finger domain (Yu et al., 2023). WRKY transcription factors activate or suppress the transcription of downstream target genes by binding to a specific *cis*-element W-box ((T) TGAC (C/T)) in downstream DNA sequences, thereby conferring specific functions on plants (Zhao et al., 2025).WRKY transcription factors in Arabidopsis, soybean, corn, poplar, potato, wheat, tomato, and Tamarix are involved in signal transduction in response to Cd stress (Wu et al., 2022; Khan et al., 2023; Liu et al., 2023; Wu et al., 2023; Xian et al., 2023). As reported, AtWRKY12 inhibits *GSH1* expression by binding to the W-box element of the *GSH1* promoter, thereby weakening tolerance to Cd (Han et al., 2019). AtWRKY13 positively regulates the expression of PDR8 by directly binding to the promoter of PDR8, thereby reducing Cd accumulation and enhancing Cd tolerance (Sheng et al., 2019). AtIRT1 is an important Fe/Cd transporter. ATL31 ubiquitizes IRT1, causing it to be degraded, while *AtWRKY33* promotes the transcription of *ATL31*. Therefore, over-expression of *AtWRKY33* reduces Cd uptake and increases Cd tolerance (Zhang et al., 2023). *WRKY45* regulates dark induced leaf senescence in Arabidopsis by regulating the expression of downstream genes (Chen et al., 2017; Barrero et al., 2022) and immune resistance in rice (SHIMONO et al., 2012; Goto et al., 2015). Arabidopsis *WRKY45* promotes transcription of the phosphate transporter *Pht1* by binding to the W-box cis element (TTGACC/T) of the phosphate transporter *Pht1;1* promoter, thereby increasing phosphorus uptake by roots (Wang et al., 2014). Our previous study found that *WRKY45* directly activates the expression of genes *PCS1* and *PCS2* that encode PC biosynthetic enzymes, thereby increasing PC content and enhancing the detoxification of Cd (Li et al., 2023). However, the transcription factors that modulate the transcription of AtWRKY45 under Cd stress are unclear.

Sulfur (S) is one of essential nutrient elements for plant growth and development. After sulfate (SO_4_^2-^) is absorbed by plants, it can gradually synthesize a variety of mercapto (-SH)-containing compounds, such as Cys, Met, GSH, and PC, which play an important role in the detoxification of heavy metals in plants (Takahashi et al., 2023). *Low Sulfate Upregulated* (*LSU*) gene family includes four genes, namely *LSU1*, *LSU2*, *LSU3* and *LSU4. LSUs* encode short peptides of about 100 amino acids (Uribe et al., 2022). It is currently known that Arabidopsis *LSU1* protein has a certain function in plant detoxification of Cd. Under Cd stress, EIN3 negatively regulates the transcription of *XTH33* and *LSU1*, and negatively regulates the growth of Arabidopsis roots. In addition, overexpressing *LSU1* in the context of *EIN3* overload (EINOX) can restore the Cd sensitivity of EIN3OX (Kong et al., 2018). LSU1 and LSU2 promote sulfur assimilation and inhibit the biosynthesis of fatty glucosinolate, promote the degradation of fatty glucosinolate, thereby limiting sulfur consumption and promoting the release of sulfur metabolites such as GSH, PC and metallothionein (MT), and increasing the Cd tolerance in Arabidopsis (Li et al., 2023).

ATP sulfurylase (APS) is a key plant enzyme that uses ATP to convert inorganic SO_4_^2-^ into organic sulfuryl sulfate (Hatzfeld et al., 2000). Arabidopsis genome has four APS genes (*APS1*, *APS2*, *APS3* and *APS4)* (Wang et al., 2020). The transcription levels of *APS* genes in Arabidopsis under Cd stress increased by more than ten times compared with the control (Harada et al., 2002). Ecotopic expressing *APS* gene in Indian mustard significantly improved their tolerance to Cd (Wang et al., 2004). Interestingly, IP-MS studies have demonstrated LSU interacts with ATP sulfurylase 1 (APS1), the #英文全 称# (GRF8) transcription factor for chlorophyll biosynthesis and seedling greening, and the ABA stress response protein RAF2/SDIRIP1 (Wang et al., 2020), but the biological significance of these interactions are unknown.

In this study, we revealed that Cd stress promotes ABA biosynthesis, ABA-biosynthesis related genes are upregulated by Cd, and the expression of *ABI5* is promoted in Cd stress. In line with this, overexpression of *ABI5* mitigates Cd stress. DUAL-LUC and EMSA verified that ABI5 directly binds the *cis*-element of WRKY45 promoter, thus boosting WRKY45 transcription. LSU1 acts as downstream target of WRKY45 verified by DUAL-LUC and EMSA. Moreover, LSU1 interacts with APS1, thus stabilizing APS1 and enhancing its enzymatic activity to assimilate sulfur. In short, we found that the ABI5-WRKY45-LSU1 axis confer the tolerance of Arabidopsis to Cd.

## 2 Materials and methods

### 2.1 Plant materials and growth conditions

Arabidopsis wild-type Col-0 was used as control genotype. The *abi5-8* (SALK_013163), *aps1-1* (SALK_046518C) and *aps1-2* (SALK_107743C) mutants were purchased from *Arabidopsis thaliana* Biological Resource Center (www.arabidopsis.org). *lsu1-1* and *lsu1-2* mutants were created via CRISPR-cas9 gene editing.

In addition, *LSU1*OX-1, *LSU1*OX-3, *APS1*OX-2, *APS1*OX-9, and LSU-GUS were single copy inserted T3 homozygous materials. *aps1-1 LSU1*OX-3 line was generated via the crossing of *aps1-1* (SALK_046518C) and *LSU1* overexpressing line.

The Arabidopsis seeds were sterilized with 75% alcohol for 1 minutes, then soaked in 95% alcohol for 30 seconds (s) and dried. After sterilization, the seeds were sown on 1/2MS medium with 0.8% sucrose and 1% agarose, and the plates were kept in the dark at 4 ℃ for 2 days for stratification treatment. The plates were then placed in a growth chamber with a thermal cycle of 22 ℃ / 20 ℃, a photoperiod of 16 hours of light / 8 hours of darkness, and a light intensity of 100 µmol^-2^s^-1^,as described (Li et al.,2023). For Cd treatment, 1/2 MS media containing 0.8% sucrose, 1% Phytagel (Sigma) and CdCl_2_ as indicated concentration were prepared and sterilized at high temperature.

#### Hydroponic culture

After the seeds were germinated and grown in 1/2MS medium for 48 hours, the seedlings were transplanted into a hydroponic device (Li et al.,2023) and grown under 100 µmol^-2^s^-1^ photosynthetically active radiation in a 16-hour light / 8-hour dark cycle. For Cd treatment, 20 μM cadmium chloride was added to the normal hydroponic nutrient solution.

### 2.2 CRISPR/Cas9 mediated gene editing to create *lsu1* mutants

In order to produce *lsu1* mutants, we used an egg-specific promoter-controlled CRISPR/Cas9 system (Wang et al., 2015). Two 19 bp single channel RNA (sgRNAs) (Xie et al., 2017) were designed for the target gene of each exon, and the target sequence was cloned into pHEE401 vector (Wang et al., 2015). The resulting plasmid was transferred into Agrobacterium tumefaciens strain GV3101, and then transformed into plants. T1 seeds were collected and spread on 1/2MS medium containing 50 mg L^-1^ hygromycin, and resistant T1 seedlings were transplanted into soil. In order to analyze the mutations in *LSU1*, genomic DNA, PCR was extracted from positive T1 plants to amplify the target site DNA containing *LSU1*. The type of mutation was identified by sequencing. To screen T_2_ plants without Cas9, two pairs of primers were used: zCas9-IDF3-2 (located in zCas9) + rbcS-E9t-IDR (located in rbcS-E9 Terminator) and zCas9-IDF3-2 + rbcS-E9tIDR2 (located in rbcS-E9 Terminator to amplify Cas9 fragments (Wang et al., 2015). The genomic DNA of Col-0 was used as negative control, and the genomic DNA from T1 transgenic Arabidopsis plants was used as positive control.

### 2.3 Recombinant DNA construction and genetic manipulation

The coding region of *Arabidopsis thaliana* LSU1 gene was amplified from cDNAs and cloned into pDONR 207 by Gateway amplification. Then the recombinant plasmid was cloned into pMDC32 vector to construct overexpression vector. In addition, the gene coding regions of *APS1* or *WRKY45* were cloned into the binary vector pCAMBIA1300 with constitutive 35S promoter to drive the expression of *APS1* or *WRKY45*. The sequence information of all genes comes from TAIR (*Arabidopsis thaliana* information resource www.arabidopsis.org). pMDC32-*LSU1*-OX and pCAMBIA1300-*APS1*-OX were introduced into Agrobacterium tumefaciens strain GV3101 by freeze-thaw method, and Col-0 were transformed with flower dipping method (Clough and Bent,1998). The transgenic lines used in this study are T_3_ homozygous plants with single copy insertion through chi-square test. In addition, the plasmid containing *LSU1* open reading frame driven by 35S promoter was transformed into *wrky45* mutant (GK-684G12) plant to produce *wrky45 LSU1*-OX, and the plasmid driven by 35S promoter *WRKY45* open reading frame was transformed into *abi5-8* (SALK_013163) plant to produce *abi5-8 WRKY45*-OX. The sequence of primer pairs for vector construction and sequencing was shown in supplementary Table S1.

### 2.4 GUS activity detection

The 2000 bp fragment of *LSU1* promoter region of *Arabidopsis thaliana* was amplified from Arabidopsis genomic DNA and cloned into pDONR207 by Gateway amplification. Then the recombinant plasmid was cloned into pMDC162 vector to construct *LSU1*pro-GUS vector. Transgenic Arabidopsis seeds germinated on 1/2 MS Agar medium and transferred to 1/2 MS Agar medium with or without 75 μM CdCl_2_ on the third day after germination. After transferring the seedlings as described, GUS staining was performed at the 6th hour of Cd treatment (Li et al., 2023).

### 2.5 Measurement of primary root length and fresh weight

After capturing the images of seedlings with a digital camera Nikon D300s, the primary root length was quantified by ImageJ (Li et al., 2023). The fresh weight of Arabidopsis was measured as described (Li et al., 2023). Statistical analysis was carried out based on three independent biological experiments.

### 2.6 RNA extraction and quantitative RT-PCR analysis

The total RNA from *Arabidopsis thaliana* seedlings was extracted and purified according to the previous method, and the purity was evaluated by A260/A280 ratio using nano-drop spectrophotometer (Thermo, USA). Use reverse transcription kit (ABclonal, China, rk 20429 https://ABclonal.com.cn/) transcribes the total RNA into cDNA. The cDNA is used for quantitative RT-PCR (qRT-PCR) analysis of SYBR Green monitoring on 7500 real-time PCR system (Thermo, USA). The primer pairs for qRT-PCR analysis of *WRKY45*, *ABI5*, *LSU1*, *LSU2*, *LSU3*, *LSU4*, *APS1*, *APS2*, *APS3*, *APS4*, *PCS1*, *PCS2*, *NACED3*, *ABA2*, *AAO3*, *CYP707A1*, *CYP707A2* and housekeeping gene *ACTIN2* (AT3G18780) were listed in supplementary Table S1.

### 2.7 Determination of cadmium content in plants

*Arabidopsis thaliana* Col-0, related mutants and transgenic seedlings were cultured in total solution for 2 weeks and then treated with 20 μM CdCl_2_ for 24 hours (h). Therefore, plants were sampled and analyzed for Cd content according to the method described (Lee et al., 2007). The accumulation and transfer factors of cadmium are calculated according to the described method (Li et al., 2023).

### 2.8 Double luciferase activity

The double luciferase experiment was basically carried out and modified as described (Li et al. 2023). *LSU1* and *WRKY45* promoters were cloned into pGreen-0800LUC vector as report vectors (LSU1Pro-LUC and WRKY45Pro-LUC). The effectors 35S::*WRKY45* and 35S::*ABI5* were constructed by using pGreenII-62-SK vector. As mentioned above, the reporter plasmid and effector plasmid were transformed into GV3101 (pSoup), and then the two kinds of *Agrobacterium tumefaciens* were simultaneously transferred to *Nicotiana benthamiana* leaves for LUC detection. Double LUC was detected by double luciferase reporter gene detection kit (Chinese https://www.yeasen.com/,YeSen) test. A modular microplate multimode reader (Turner Biological system) is used to detect LUC and any signal.

### 2.9 Protein extraction and western blot analysis

Total protein is extracted as described earlier (Li et al., 2023). Arabidopsis plants were ground in liquid N_2_ and extracted with an extraction buffer (50 mM Tris pH 7.5,150 mM sodium chloride; 5mM EDTA;2mm DTT;10% glycerin; 1 mm PMSF). After centrifugation at 4 ℃ with 12,000 rpm for 10 minutes, the supernatant was loaded onto 10% SDS-PAGE and western blotting was performed using anti-GFP (M20004) or anti-actin antibody (M20009) (Abmart, Chinese http://www.ab-mart.com.cn/).

### 2.10 *In vitro* pull-down analysis

As mentioned above, the complete coding sequence of *APS1* was constructed into pGEX-6P-3 vector, and the recombinant expression vector pGEX-6P-3-*APS1* was introduced into *E.coli* BL21 strain, and the GST-APS1 protein was expressed and purified. LSU1-His protein was also expressed and purified according to the above method. For in *vitro* pull-down assay, 10 mg recombinant protein (GST-APS1) and 10 mg LSU1-His were incubated in 500 microliters of binding buffer (2 mM DTT, 10-mM MgCl2,20 mM Tris-HCl, pH 7.2) for 1 hour. Then Bever Beads^TM^ GSH magnetic beads (Beaver, 70601 https://www.beaverbio.com/) were incubated with bound buffer for 20 minutes and washed for 5 times with GSH eluent. The pulled proteins were detected by immunoblotting with anti-HIS antibody (Yesen, 30401ES, China, https://www.yeasen.com/).

### 2.11 Co-IP analysis

To detect the interaction between LSU1 and APS1, we amplified the full-length cDNAs of *LSU1* and *APS1*, and constructed pCAMBIA1300-*APS1*-GFP and pCAMBIA1300-*LSU1*-Flag vectors. The primers used are listed in supplementary Table S1. *Agrobacterium tumefaciens* containing recombinant APS1-GFP and LSU1-Flag were transferred into *Arabidopsis thaliana* respectively, and stable transgenic lines were obtained. Total proteins were extracted from transgenic leaves with buffers containing 50 mM HEPES-KOH (pH 7.5), 100 mM NaCl, 0.1%Triton 10, 5% glycerol and protease inhibitors. The mixed protein extract was incubated with Flag magnetic beads (M20118, Abmart, http://www.ab-mart.com.cn//) for 20 minutes and washed for 5 times to remove non-specific binding proteins, followed by protein elution with 3 × Flag peptides. The eluted samples were analyzed by immunoblotting with GFP antibody (M20004, Abmart) and FLAG (M20008, Abmart) antibody, respectively.

### 2.12 Electrophoresis mobility shift assay (EMSA)

We used EMSA gel shift kit (GS009, Beyotime, China https://www.beyotime.com/index.htm) to explore the binding of ABI5 with the promoter of *AtWRKY45*. The complete coding sequence of WRKY45 was amplified by primers GST_WRKY45_F and GST_WRKY45_R. The amplification product of WRKY45 was cloned into pGEX-6P-3 vector (GE Healthcare, USA) cut by BamHI.

The expression vector was introduced into *E. coli* BL21 strain. The recombinant protein GST-WRKY45 was extracted with BeverBeads^TM^ GSH magnetic beads (70601, Beaver Chinese https://www.beaverbio.com/). Similarly, all the primers *ABI5*-His-F and *ABI5*-His-R were used to amplify the complete coding sequence of *ABI5*. The amplification product of *ABI5* was cloned into pET29b vector cut by BamHI and SalI, and the vector was also introduced into *E coli* BL21 strain. The BeverBeads^TM^ His-tag protein was purified by magnetic beads (70501, Beaver Chinese https://www.beaverbio.com/). Oligonucleotide probes (*LSU1*-EMSA-bio and *WRKY45*-EMSA-bio) were used and the 5 ′ends were labeled with biotin. The sequence is shown in the Table S1.

### 2.13 Split-LUC test

Split-LUC method was used to confirm the interaction between LSU1 protein and APS1 protein. PCAMBIA1300nLUC-LSU1 and pCAMBIA1300-cLUC-APS1 were transformed into *Agrobacterium tumefaciens* GV3101 and expressed transiently in Nicotiana *benthamiana* leaves. After cultured in the growing chamber for 3 days, the leaves were sprayed with 1 mM of D-fluorescein potassium salt (40902ES ·YeSen, China, https://www.yeasen.com/). Then the 5 min was preserved in the dark, and then the luminous signal was detected by plant *in vivo* imaging system (Night SHADE; Berthold).

### 2.14 Analysis of PC, NPT, glutathione, and chlorophyll content

Two-week-old seedlings cultured in water for 24 hours were treated with 20 Mm CdCl_2_, and then the contents of PC, NPT and GSH were analyzed and determined as described (Li *et al*. 2023). The seedlings were cultured on a 1/2 MS medium with or without 75 μM CdCl_2_ for 10 days, and then the chlorophyll content was analyzed based on an open method (Nam *et al*., 2021).

### 2.15 Quantification of ATP sulfurylase activity

Two-week-old seedlings cultured in water for 6 hours and 24 hours were treated with 20 μM CdCl_2_, and then ATP sulfurylase activity was analyzed and determined as described (Adams et al., 1968).

### 2.16 RNA-seq analysis

We exploited RNA-Seq is completed by Novogene Bioinformatics Technology Co (Shanghai, China). Total RNA was extracted with general plant total RNA extraction kit (BioTeke, Beijing, China), and then treated with Turbo DNA-free kit (Ambion, USA). Transcriptome sequencing and data analysis were performed as previously described (Li et al., 2024). Gene expression levels were quantified as fragments per kilobase per million mapped reads (FPKM). The transcriptome data has been deposited in the National Center for Biotechnology Information (NCBI) under accession number PRJNA1374680. A hierarchical clustering analysis of the DEGs was performed to explore gene expression patterns. The GO enrichment of the DEGs was estimated using the DESeq R package based on the hypergeometric distribution.

## 3. Results

### 3.1 Cd stress increases ABA biosynthesis

Exogenous ABA can alleviate the symptoms of cadmium poisoning in plants (Hu et al., 2020). In line with this, as shown in Fig.S1, ABA application effectively increased chlorophyll content and fresh weight of Arabidopsis seedling. Compared with control (no Cd treatment), the primary root (PR) length of Col-0 was decreased by 54.7% under 75 μM Cd treatment (Fig. S1 B), fresh weight (FW) was reduced by 67.9%(Fig. S1 C), chlorophyll content was decreased by 63.6%(Fig. S1 D); When 0.5μM ABA was added, the PR length of Col-0 was significantly reduced by 28.9% compared to 75μ M Cd treatment (Fig.S1 B), but FW and chlorophyll content were increased significantly by 11.1% (Fig.S1 C) and by 17.6%(Fig.S1 D), respectively. Application of 2μM ABA to Cd reduced the PR by 30.4% in contrast to the control (Fig.S1 B), and the FW and chlorophyll content were decreased significantly by 17.2% (Fig.S1 C), and by 27.6% (Fig.S1 D), respectively. However, 4 μM ABA significantly reduced the PR length of Col-0 by 50.6% (Fig.S1 B), FW and chlorophyll content were decreased by 30.4% (Fig.S1 C) and 30.6% (Fig.S1 D). Our results indicates that ABA can effectively mitigate Cd stress in Arabidopsis, especially improved chlorophyll content and fresh weight.

We suspected that Cd stress increases the biosynthesis of ABA in Arabidopsis. To verify that, we analyzed the transcripts of ABA synthesis-related genes (*NCED3, ABA2, AAO3*) and degradation related genes (*CYP707A1, CYP702A2*) in cadmium-treated Arabidopsis plants. As shown in Fig.S1, the transcripts of *NCED3* were up-regulated continuously within 6 hours of cadmium stress, and peaks at 6th hour (Fig.S1 E). The transcripts of *ABA2* and *AAO3* were up-regulated and then decreased within 6 hours of cadmium stress, and peaked at third hour, which were 25.7 and 14.8 times higher than those at 0 hours, respectively (Fig.S1 F G). On the contrary, the transcripts of *CYP707A1* and *CYP702A2* were decreased continuously within 6 hours of cadmium stress and reached the lowest value at 6 hours, which were 0.22 and 0.41 times higher than those of 0 hours, respectively (Fig.S1 H I). Consistent with this finding, ABA level was also significantly increased after cadmium treatment, which increased by 2.41 and 2.59 folds after 6 h and 24 h of cadmium treatment, respectively (Fig.1 A). Based on our results, we thus hold that Cd stress stimulates the ABA biosynthesis.

**Fig. 1.**
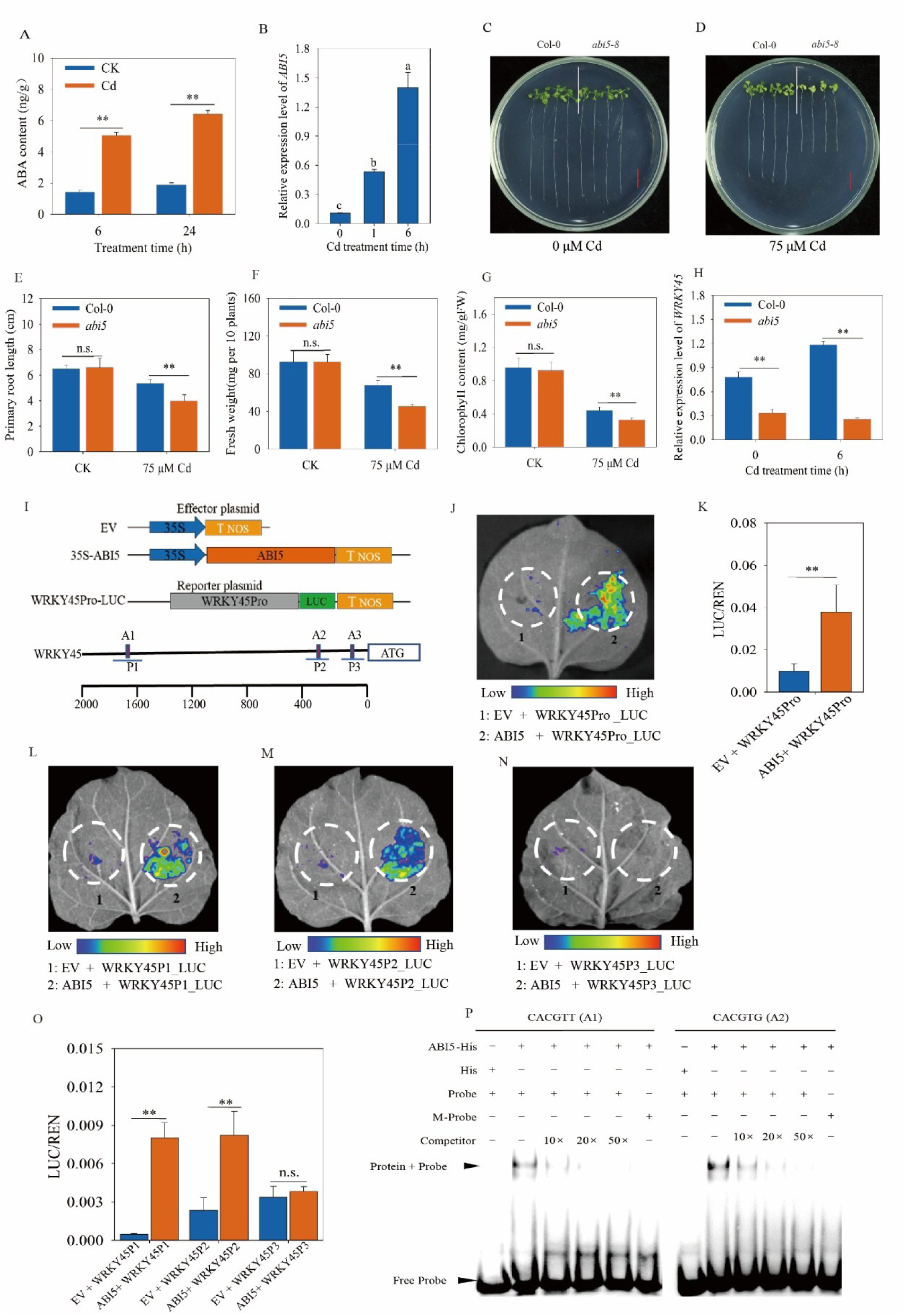
ABI5 promotes *WRKY45* transcription by binding to the ABRE element of the *WRKY45* promoter and thus positively regulates Cd tolerance in Arabidopsis. (A) The relative expression level of *ABI5* in Arabidopsis. qRT-PCR analysis of transcript levels of *ABI5* in Arabidopsis at different time point after Cd stress. (B) Contents of endogenous ABA in Arabidopsis treated with Cd for 6 h and 24 h. (C, D) The *abi5-8* mutant plants are more sensitive to Cd. Two-day old plants grown on half strength Murashige and Skoogs (1/2 MS) medium were transferred to 1/2 MS medium without or with 75 μM CdCl_2_. Photographs were taken 10 days after the transfer. Scale bar, 1 cm. (E, F, G) Primary root length (E), fresh weight (F) and chlorophyll (G) were measured. Three independent experiments were done with similar results, four plants per genotype from one plate were measured for each repeat. (H) The expression of *WRKY45* in Col-0, and *abi5* lines by qRT-PCR. Two-week-old seedlings were treated with 20 µM CdCl_2_ for 0 h and 6 h. (I) Schematics of all constructs used for transient expression assays in *Nicotiana benthamiana* leaves. The WRKY45 promoter was fused to the luciferase (LUC) reporter gene. 35S::*ABI5* acted as an effector, and the empty vector (EV) acted as the negative control; and ABI5-binding site (ABRE, CACGT[T/G]) prediction within 2000 bp of *WRKY45* promoters. The WRKY45 promoter was divided into four segments, namely *WRKY45* probe 1 (*WRKY45*-P1), *WRKY45*-P2, and *WRKY45*-P3. (J) LUC fluorescence imaging shows expressions of *WRKY45*Pro after co-expression with *ABI5*. (K) The relative LUC/REN reporter activity was quantified. (L-N) LUC fluorescence imaging shows that *ABI5* activates *WRKY45*-P1 (L) and *WRKY45*-P2 (M) but not *WRKY45*-P3(N), respectively. (O) The relative LUC/REN reporter activity was quantified. *WRKY45*P1-LUC, *WRKY45*P2-LUC and *WRKY45*P3-LUC, was co-transformed with 35S:: WRKY45 into *N.benthamiana*. (P) Electrophoretic mobility shift assay (EMSA) shows that ABI5 bounds to ABRE1(A1) in *WRKY45*-P1 probe and ABRE2(A2) in *WRKY45*-P2 probe, but not mutated ABRE1 (mABRE1) and ABRE2 (ABRE2). Data are means ± SE (n = 4). Different letters indicate significant difference at *P <* 0.05 (one-way ANOVA with Turkey’s test), and the asterisks indicate significant differences from the control (*, *P <* 0.05, **, *P <* 0.01; Student’s *t*-test).

Given that ABI5 is the hub transcription factor in ABA signaling (Hu et al., 2020), we also explored the its transcripts in Cd stress. As expected, the transcripts of ABI5 were up-regulated within 6 hours of cadmium stress, and the highest expression was reached at 6 hours, which was 13.1 times higher than that of 0 hours under cadmium stress. (Fig.1 B). This indicates that ABI5 is involved in Cd tolerance. Moreover, the expression of *ABI5* was up-regulated within 24 hours after exogenous ABA treatment, and reached the peak at 6 hours of treatment, which was 12.5 times higher than that of ABA treatment for 0 hours (Fig.S2 A). To clarify the roles of *ABI5* in coping with Cd stress, we firstly check the sensitivity of *abi5-8* mutant to Cd. Compared with Col-0, no significant growth and developmental defects were observed in *abi5-8* at seedling stage when growing in 1/2 MS media (Fig. 1C). However, *abi5-8* mutants are more sensitive to Cd than Col-0 (Fig. 1D). We therefore quantified the PR length (Fig. 1E), FW (Fig. 1F) and chlorophyll content (Fig. 1G). Under the 0 μM CdCl_2_ no significant difference was observed between Col-0 and *abi5-8* plants. When exposed to 75 μM CdCl_2_, *abi5-8* mutant plants showed a 30.1% reduction in PR length compared to Col-0, along with a 25.8% decrease in FW and a 10.1% decline in chlorophyll content (Fig. 1E F G). In short, these results support that the ABI5 positively regulate Cd tolerance in Arabidopsis.

Our previous studies have shown that *WRKY45* increases Cd tolerance in Arabidopsis by positively regulating the expression of *PCS1* and *PCS2* (Li et al.,2023). We were curious about the responses of *WRKY45* to ABA. The addition of ABA increased the transcription level of *WRKY45*. The transcription level of *WRKY45* peaked at the 24^th^ ABA treatment, which was as 10.8 times as that of the ABA treatment 0 h (Fig.S2 B). To link ABI5 with the expression of *WRKY45* under Cd stress, we detected the transcripts of *WRKY45* at Col-0 and *abi5-8*. The transcript level of *WRKY45* in Col-0 was much higher than that in *abi5-8* at 0 h and 6 h of Cd treatment, which increased by 2.01 and 3.67 times, respectively (Fig. 1H). These results suggest that *ABI5* appears to promotes *WRKY45* expression in response to Cd stress.

Arabidopsis *ABI5* regulates the expression of downstream genes by binding ABRE *cis*-element (CACGT[T/G]) (Wang et al.,2019). We found three ABRE elements in *WRKY45* promoter (Fig. 1 I). This hint at that ABI5 may directly bind to the ABRE element in the promoter region of *WRKY45* to promote its transcription. To verify that, we fused the promoter of *WRKY45* with luciferase reporter gene (Fig. 1I). We then constructed 35S promoter-driven effect vector (35S::*ABI5*) and co-transformed the reporter vector into *Nicotiana benthamiana* as described. Fluorescence imaging showed that *ABI5* promoted the expression of luciferase driven by *WRKY45* promoter (WRKY45Pro) (Fig. 1J). In contrast to the control vector, the expression of *ABI5* significantly increased the LUC/REN ratio (Fig. 1K), suggesting that ABI5 binds the promoter of *WRKY45*.

To test whether ABI5 binds to the ABRE element of *WRKY45* promoter, three fragments of *WRKY45* promoter region were cloned, each fragment containing ABRE *cis-*element and named WRKY45P1(−1759-2036bp), WRKY45P2(−192-597bp) and WRKY45P3 (−8-145bp) based on the upstream position of the initiation codon ATG. Then, these fragments were fused with luciferase reporter vector and expressed in *Nicotiana benthamiana* together with *ABI5* effect vector driven by 35S promoter. Fluorescence imaging showed that *ABI5* specifically enhanced the expression of luciferase reporter vector driven by P2 and P3 fragments of *WRKY45* promoter, but had no effect on *WRKY45P*1 fragment (Fig. 1L M N). Compared with the control vector, *ABI5* significantly enhanced the LUC/REN ratio in *WRKY45*P2 and *WRKY45*P3 (Fig. 1O).

In addition, EMSA experiment confirmed that ABI5 can directly bind ABRE1(A1) probe located in P1 segment of WRKY45 promoter and ABRE2(A2) probe located in P2 segment. After adding 20 times of competitive cold probe, the binding of ABI5 to ABRE1(A1) was almost completely inhibited, while the binding of ABI5 to ABRE2(A2) needed 50 times of competitive cold probe to be inhibited. This suggests that the binding strength of ABI5 with A2 is stronger. Furthermore, ABI5 could not bind to the mutated ABRE probe mabre (“GAACG” or “ATCTA”) (Fig. 1P). These results proved that ABI5 regulates the transcription of *WRKY45* by directly binding the ABRE element in the promoter region of *WRKY45 in vitro*.

To clarify the genetic interaction between *ABI5* and *WRKY45*, we overexpressed *WRKY45* in the abi5-8 mutant and obtained two homozygous lines, *abi5*/WRKY45OX-6 and a*bi5*/WRKY45OX-13, which were screened by qRT-PCR (Fig. 2A). In 1/2MS medium without cadmium, there was no significant difference in growth between *abi5*/WRKY45OX-6, *abi5*/WRKY45OX-13, Col-0 and the *abi5-8* mutant (Fig. 2B). However, under 75 μM Cd²⁺ stress, compared with the *abi5-8* mutant and Col-0, the *abi5*/WRKY45OX-6 and *abi5*/WRKY45OX-13 lines exhibited Cd tolerance in terms of PR length (Fig. 2C), FW (Fig. 2D), and chlorophyll content (Fig. 2E). Therefore, we conclude that *WRKY45* functions downstream of *ABI5* and is involved in regulating plant response to Cd stress.

**Fig. 2.**
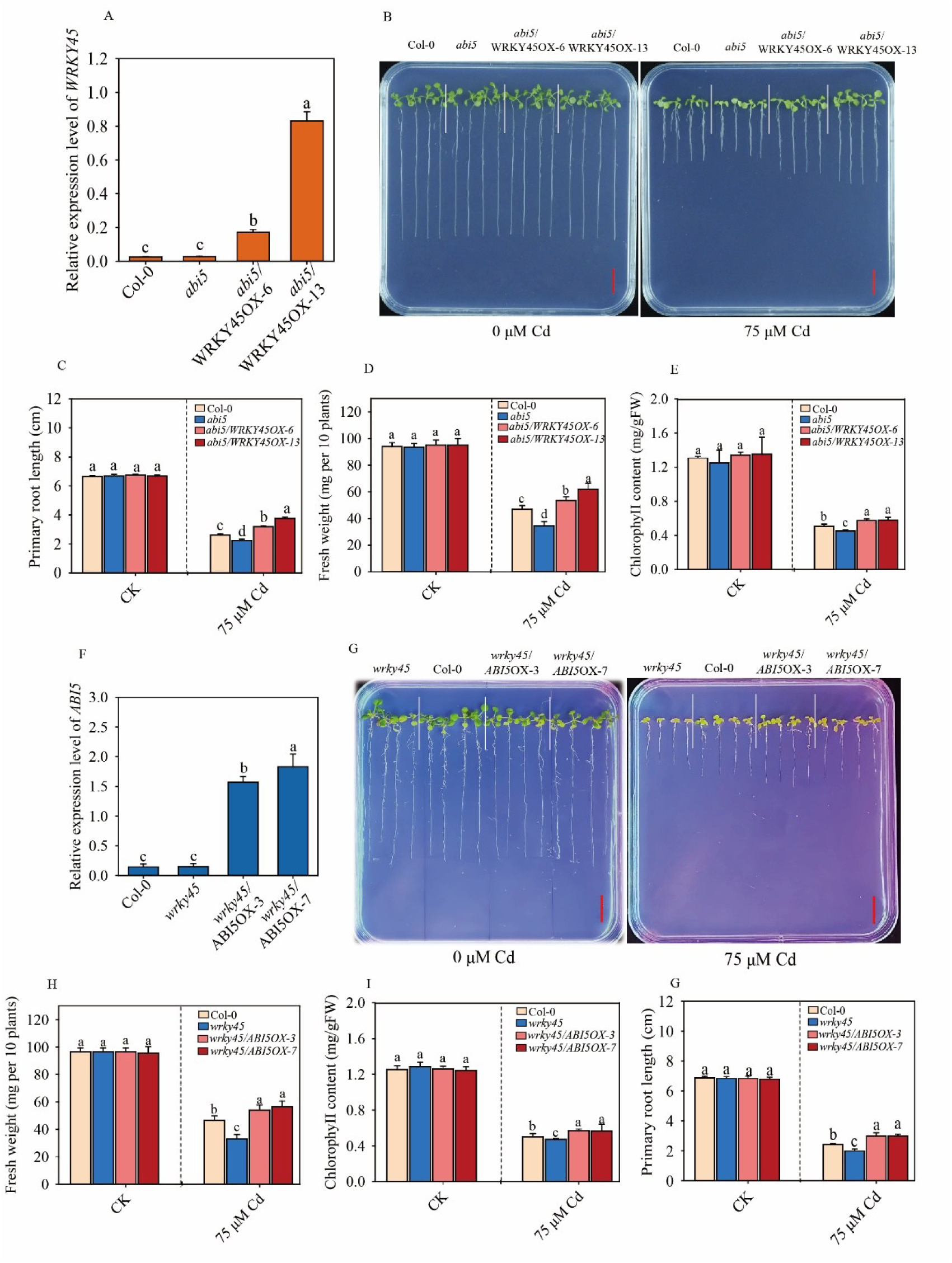
*ABI5* positively regulates Cd detoxification and acts as upstream of *WRKY45*. (A) Expression of WRKY45 of *abi5-8*, wild type Col-0, *abi5*/WRKY45OX-6, and abi5/WRKY45OX-13 lines. (B) Growth of *abi5-8*, wild type Col-0, *abi5*/WRKY45OX-6, and abi5/WRKY45OX-13 lines under Cd stress. Two-day-old plants grown on 1/2 MS media were transferred to 1/2 MS media containing 0 or 75 μM CdCl_2,_ respectively. Photographs were taken 10 days after treatment. Scale bar, 10 mm. Primary root length (C), fresh weight (D) and chlorophyll (E) of plants were measured. (F) Expression of *ABI5* of *wrky45,* Col-0, wrky45/ABI5OX-3, and wrky45/ABI5OX-7 lines. (G) Growth of *wrky45*, Col-0, *wrky45*/ABI5OX-3, and *wrky45*/ABI5OX-7 lines under Cd stress. Photographs were taken 10 days after the treatment. Scale bar, 10 mm. Primary root length (H), fresh weight (I) and chlorophyll (J) of plants were measured. Three independent experiments were done with similar results, each with three biologicals repeats. Four plants per genotype from one plate were measured for each repeat. Data are presented as means ± SE (n = 4). Different letters indicate significant difference at *P <* 0.05 level (one-way ANOVA with Turkey’s test).

In addition, we overexpressed *ABI5* in the *wrky45* mutant and obtained two homozygous lines, *wrky45*/ABI5OX-3 and *wrky45*/ABI5OX-7, through qRT-PCR screening (Fig. 2F). In 1/2MS medium without Cd, there was no significant difference in growth between *wrky45*/ABI5OX-3, *wrky45*/ABI5OX-7, Col-0 and the *wrky45* mutant (Fig. 2G). However, under 75 μM Cd²⁺ stress, compared with the wrky45 mutant and Col-0 wild type, the wrky45/ABI5OX-3 and wrky45/ABI5OX-7 lines showed Cd tolerance in terms of PR length (Fig. 2H), FW (Fig. 2I), and chlorophyll content (Fig. 2G). It is possible that *ABI5* has other downstream genes besides those downstream of *WRKY45*.

### 3.2 *WRKY45* positively regulates the transcription of *LSU1*

Our previous studies have shown that overexpression of *WRKY45* increases the content of phytochelatin (PC) and non-protein thiol (NPT) in plant. Consistently, the contents of PC and NPT in *wrky45* mutants are decreased significantly (Li et al., 2023). Apart from *PCS1* and *PCS2*, other targets of WRKY45 remain elusive. To mine out other target genes, we thus exploited transcriptome sequencing to look for other downstream genes of WRKY45 involved in Cd tolerance. The transcriptomes of root and leaf samples from Col-0, WRKY45-OX-15, *wrky45* were subjected to sequencing after treatment with 20 μM Cd in hydroponic culture for six hours. The VENN map showed that there were 150 and 169 overlapping genes between WRKY45-OX vs Col-0 and Cd vs CK in roots and leaves; 130 and 66 overlapping differentially expressed genes (DEGs) between *wrky45* vs Col-0 and Cd vs CK in roots and leaves (Fig. 3A, Fig. S3A); 323 and 74 overlapping genes between *wrky45* vs Col-0 and WRKY45-OX vs Col-0 in roots and leaves of *Arabidopsis thaliana* (Fig. 3A, Fig. S3A). *WRKK45* overexpressing and *wrky45* mutation have a lot of overlapping DEGs regulated by Cd stress, respectively. We summarized the above DEGs and analyzed them by GO enrichment, and found that the sulfur (S) compound metabolic process (GO:0006790) pathway was enriched in Arabidopsis roots and leaves (Fig. 3B, Fig. S3B).

**Fig. 3.**
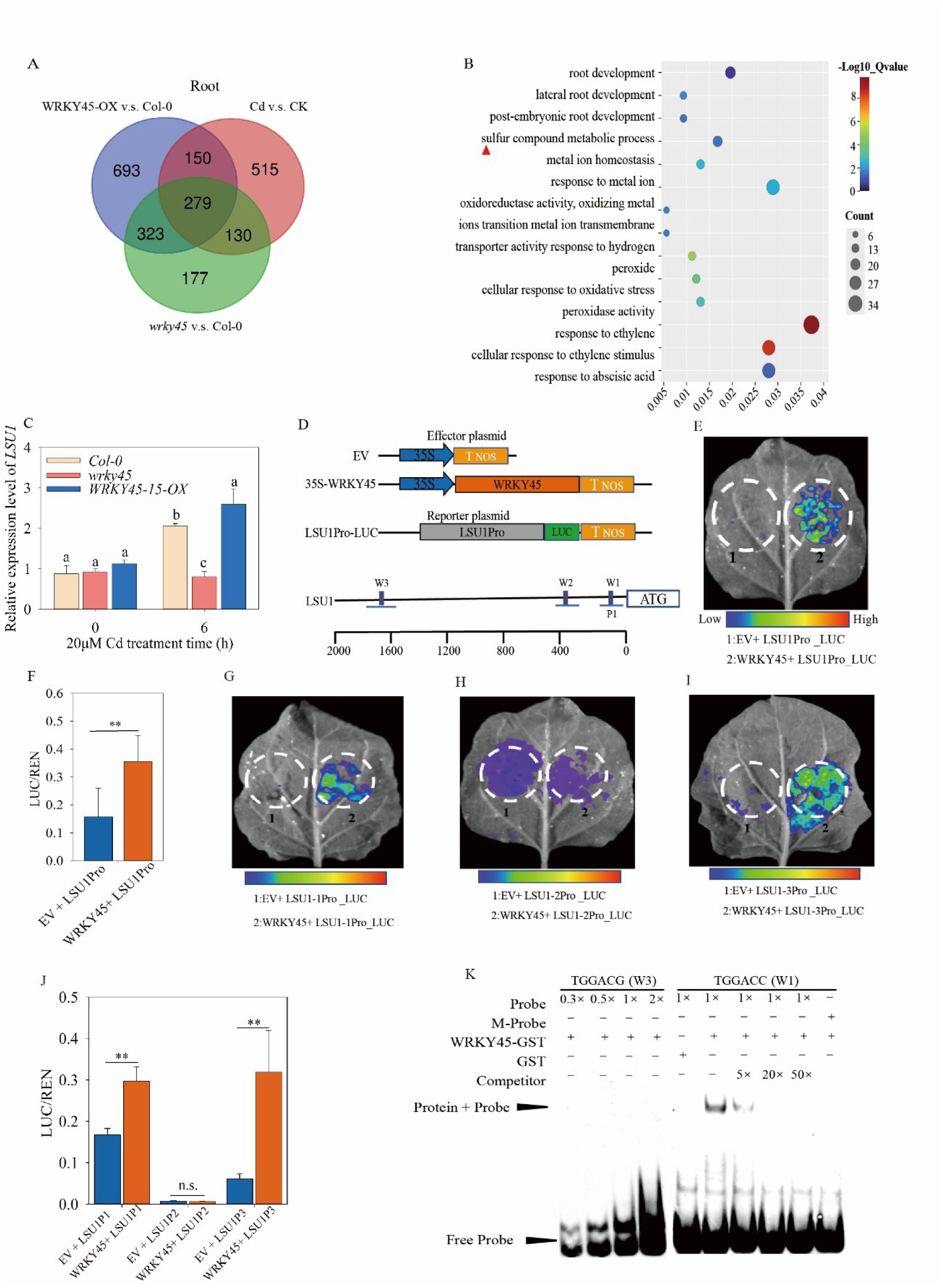
WRKY45 promotes *LSU1* transcription by binding to the W-box element of the *LSU1* promoter and thus positively regulates Cd tolerance in Arabidopsis. (A) Venn diagram illustrating the differentially expressed genes (DEGs) shared between Col-0, *wrky45* and WRKY45-OX-15 roots after 6 hours of Cd exposure. (B) GO term enrichment analysis of DEGs in the biological process in Col-0, *wrky45* and WRKY45-OX-15 roots after 6 hours of Cd exposure. (C) The expression of *LSU1* in Col-0, *wrky45*, and *WRKY45-OX* lines by qRT-PCR. Hydroponics two-week-old seedlings were treated with 20 µM CdCl_2_ for 0 h and 6 h. The relative expression was normalized against *ACTIN2* (*AT3G18780*). (D) Schematics of all constructs used for transient expression assays in *Nicotiana benthamiana* leaves; and WRKY45-binding site (W-box, TTGAC[C/T]) prediction within 2000 bp of *LSU1* promoters. The *LSU1* promoter was divided into three segments, namely *LSU1* probe 1 (LSU1-P1), LSU1-P2, and LSU1-P3. (E) LUC fluorescence imaging shows expressions of *LSU1*Pro after co-expression with *WRKY45*. (F) The relative LUC/REN reporter activity was quantified. (G-I) LUC fluorescence imaging shows that WRKY45 activates LSU1-P1 (G) and LSU1-P3 (I) but not LSU1-P2(H), respectively. (J) The relative LUC/REN reporter activity was quantified. LSU1P1-LUC, LSU1P2-LUC and LSUI1P3-LUC, was co-transformed with 35S: WRKY45 into *N.benthamiana,* respectively. (K) Electrophoretic mobility shift assay (EMSA) shows that WRKY45 protein bounds to W-box1(W1) in LSU1-P1 probe, but not mutated W-box1 (mW-box1). Data are presented as means ± SE (n = 4). The asterisks indicate significant differences from the control (*, *P <* 0.05, **, *P <* 0.01; Student’s *t*-test). n.s., not significant. Different letters indicate significant difference at *P* < 0.05 level (one-way ANOVA with Turkey’s test).

As reported, LSU gene family are induced by S deficiency (Niemiro et al., 2020) and Cd stress (Li et al., 2023). We noticed that LSU1 was up-regulated genes in WRKY45-OX background in contrast to Col-0 (Table S2).

We first detected the transcript level of *LSU1* at Col-0, *wrky45* and WRKY45-OX-15 levels. Of note, at 0 h of Cd treatment, the expression level of *LSU1* was almost indistinguishable in Col-0, *wrky45* and WRKY45OX-15 roots; however, after 6 hours of Cd treatment, the expression level of *LSU1* increased in all three genotypes, and the highest abundance was observed in WRKY45OX-15 roots, followed by Col-0 and *wrky45* (Fig. 3C). Our results show that *WKKY45* promotes the expression of *LSU1* in response to Cd stress.

To verify that LSU1 is the target gene of WRKY45 at transcription level, we cloned the native promoters of *LSU1* and fused them with the luciferase reporter gene (Fig. 3D). Then, we co-transformed *Nicotiana benthamiana* with a plasmid expressing *WRKY45* driven by 35s promoter. Fluorescence imaging showed that *WRKY45* promoted LSU1Pro-LUC (Fig. 3E), and the expression of *WRKY45* in *LSU1* promoter increased LUC/REN ratio to a higher extent than that of empty vector (EV) (Fig. 3F). Moreover, we found that that the promoter region of *LSU1* contained W-box cis elements (Fig. 3D). We assumed that WRKY45 binds directly to these W-boxes and affects *LSU1* expression. To test this, three fragments of the promoter region of *LSU1* were cloned, each containing W-box and named as LSU1P1 (0 - −234bp), LSU1P2 (−506 - −581bp) and LSU1P3 (−1825 - −1967bp) upstream of the initiation codon ATG (Fig. 3D). Fluorescence imaging revealed that *WRKY45* specifically enhanced the expression of luciferase reporter gene driven by P1 and P3 fragments of *LSU1* promoter, but had no effect on P2 (Fig. 3G H I). Compared with the empty vector, the overexpression of *WRKY45* significantly enhanced the luciferase activity (Fig. 3J). In addition, our EMSA assay confirmed that the direct binding of WRKY45 to the biotin-labeled W-box1 probe located in the P1 region of the *LSU1* promoter was almost completely inhibited by the addition of 20-fold excess of unlabeled competitive no-cold probe. Of note, WRKY45 could not bind to the mutated W-box probe mW-box (“CGCGGA”) (Fig. 3K). These results confirmed that WRKY45 regulated the transcription of *LSU1* by directly binding to W-box1 in the promoter region of *LSU1 in vitro*. On the other hand, we tried to confirm the interaction between W-box3 motifs in *LSU1* and WRKY45 by EMSA analysis, but failed.

### 3.3 *LSU1* positively regulates cadmium tolerance through glutathione-dependent PC synthesis pathway

We studied the transcripts of *LSU1* under Cd stress by qRT-PCR technique. The transcription level of *LSU1* in roots and leaves of *Arabidopsis thaliana* was enhanced by Cd stress. Compared with the control (without Cd), the expression of *LSU1* increased significantly at 6 and 24 hours after Cd stress (Fig. 4A B). In roots, when treated with 6 h Cd, the abundance of *LSU1* was the highest, which increased by 2.6 times (Fig. 4A), while in leaves, when treated with 24h Cd, the abundance of *LSU1* was the highest, which increased by 1.9 times. Our results confirmed that *LSU1* was induced by cadmium stress.

**Fig. 4.**
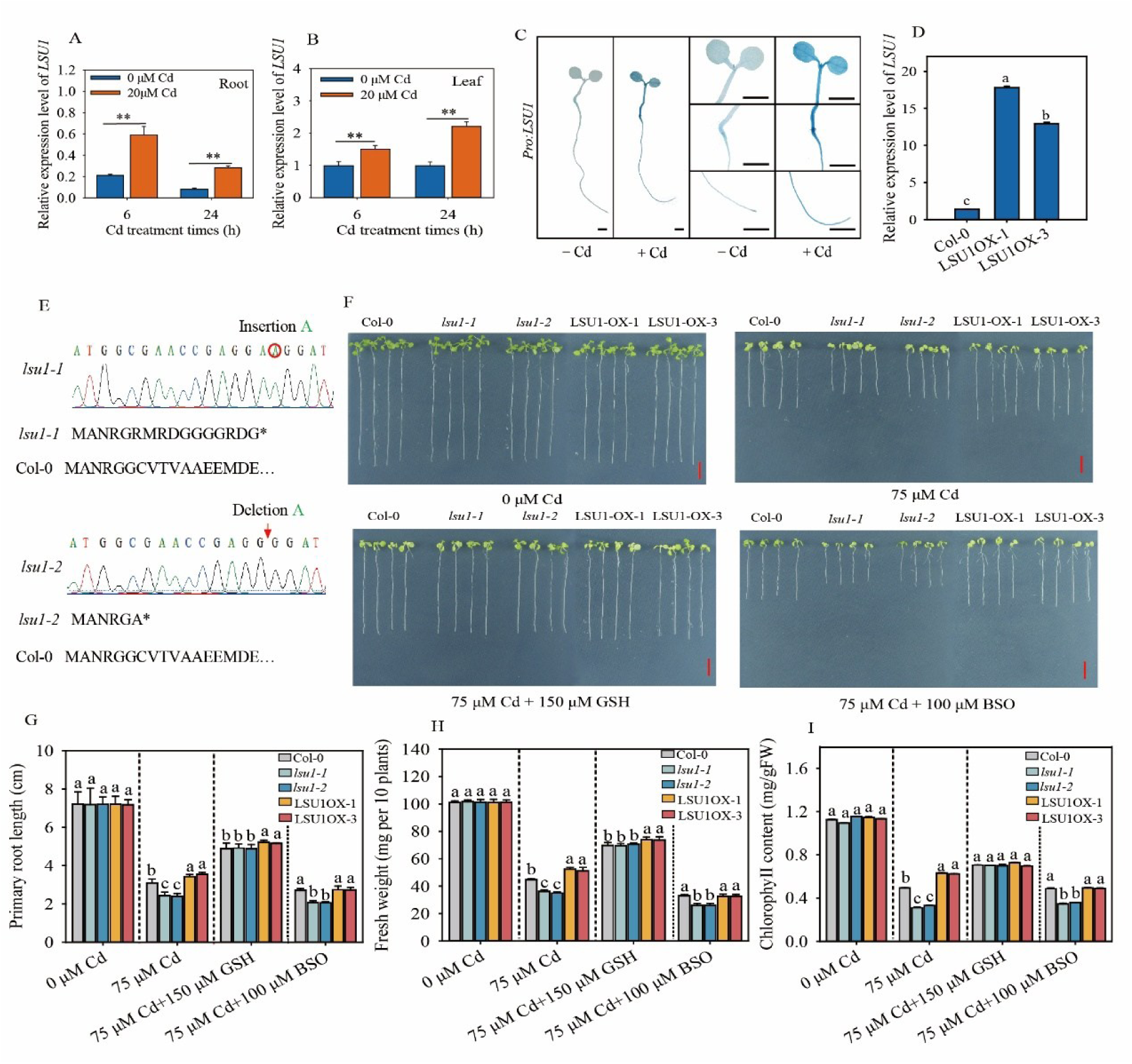
*LSU1* upregulated expression induced by Cd stress and positively regulated the detoxification of Cd in Arabidopsis. (A, B) The relative expression levels of *LSU1* in leaves and roots of Arabidops were analyzed by qRT-PCR. The asterisks indicate significant differences from the control (*P <* 0.01, Student’s test). (C) Cd stress promoted the activity of *LSU1* promoter. ProWRKYLSU1:: GUS seedlings were stained as described. The seeds of Arabidopsis germinated on ½ agar medium and transferred to solid agar media with or without 75 μM CdCl_2_ 3 days after germination. 24 hours after transplanting, the results were as follows: cotyledon, scale bar, 2 mm; root connection part, scale bar, 2 mm; primary root, scale bar,2 mm. (D) The expression level of *LSU1* in *LSU1* overexpression materials. The relative expression level of *LSU1* was normalized with *ACTIN2* (*AT3G18780*). (E) Sequencing results of *lsu1-1* and *lsu1-2* mutants. (F-I) LSU1-mediated Cd tolerance depended on endogenous glutathione biosynthesis. (F)Arabidopsis thaliana was grown on 1/2MS media containing 0 μM Cd, 75 μM Cd, 75 μM Cd+150 μM GSH, 75 μM Cd + 100 μM BSO, respectively. The seedlings were grown on half-strength MS media for 2 days, and then treated with 0 μM Cd, 75 μM Cd, 75 μM Cd+150 μM GSH, 75 μM Cd + 100 μM BSO for 10 days. Scale bar, 1 cm. The primary root length (G), fresh weight (H) and chlorophyll content (I) of Col-0, *lsu1-1*, *lsu1-2*, LSU1OX-1 and LSU1OX-3 were measured. Three independent experiments were done with similar results. Four plants per genotype from one plate were measured for each repeat. Data present the means ± SE (n = 4). Different letters indicate significant difference at *P <* 0.05 level (one-way ANOVA with Turkey’s test).

As reported, *LSU2, LSU3*, *LSU4* are homologue of *LSU1* (Sirko et al., 2015). We subsequently revealed that the expression of *LSU2*, *LSU3* transcripts in Arabidopsis roots and leaves was the highest under Cd stress for 6 hours; in Arabidopsis roots and leaves, the expression of *LSU2* transcripts was also significantly up-regulated after 24 hours of Cd stress, while *LSU3* was significantly up-regulated in root samples under Cd for 24 hours, but not in leaf samples. The transcript of *LSU4* did not seem to be strongly induced by Cd stress. In our experiment, the up-regulated expression of *LSU4* was only detected in root samples at 24 h under Cd stress (Fig. S4A B C). This suggests that *LSU2* and *LSU3* may also play roles in Cd tolerance.

We obtained ProLSU1: GUS plants by transgenic technology and evaluated the activity of *LSU1* promoter under Cd stress. Under the control condition (-Cd), we observed the activity of *LSU1* promoter in roots, stems and leaves. It is worth noting that the activity of *LSU1* promoter in roots and leaves increased significantly under Cd stress (+Cd) (Fig. 4C). This result further supports the positive response of *LSU1* to Cd.

In order to decipher the physiological function of *LSU1* under Cd stress, we used CRISPR/Cas9-mediated gene editing technique to obtain *LSU1* gene editing mutants *lsu1-1* and *lsu1-2* (Fig. 4E). In addition, we also created an overexpression line of *LSU1* (Fig. 4D). Our sequencing method confirmed that the *lsu1-1* mutant inserted a base (A) after the 15th base after the start codon ATG, while the *lsu1-2* mutant deleted one base (A) after the 14th base after the start codon ATG (Fig. 4E). The mutations of *lsu1-1* and *lsu1-2* results in the early termination of *LSU1* translation (Fig. 4). Furthermore, we created overexpressing *LSU1* lines (LSU1OX-1 and LSU1OX-3). Compared with Col-0, the expression levels of *LSU1* were 1.9 and 8.4-fold higher in LSU1OX-1 and LSU1OX-3 materials, respectively (Fig. 4D). We thus used these materials in subsequent experiments.

Compared with Col-0, no significant growth and developmental defects were observed in *lsu1-1*, *lsu1-2*, LSU1OX-1 and LSU1OX-3 lines when grown in 1/2 MS media. However, when exposed to Cd, overexpressing *LSU1* plants showed higher Cd tolerance than Col-0, while *lsu1* mutant plants were more sensitive to Cd (Fig. 4F). We quantified the PR length (Fig. 4G), FW (Fig. 4H) and chlorophyll content (Fig. 4I). Under the condition of 0 μM CdCl_2_, no significant difference was observed between Col-0, *lsu1-1*, *lsu1-2*, LSU1OX-1 and LSU1OX-3 plants. However, when exposed to 75 μM CdCl_2_ stress, *lsu1-1* and *lsu1-2* mutant plants showed a significant decrease in PR length by 21.5% and 22.6% in contrast to with Col-0. LSU1OX-1 and LSU1OX-3 plants showed a significant increase of 19.0% and 14.9%, respectively (Fig. 4G). Under these conditions, the FW of *lsu1-1* and *lsu1-2* plants decreased by 19.0% and 21.8% respectively, while the FW of overexpressing LSU1OX-1 and LSU1OX-3 plants increased by 17.2% and 14.4%, respectively (Fig. 4H). When exposed to 75 μM CdCl_2_, the chlorophyll content in *lsu1-1* and *lsu1-2* plants was lower than that in Col-0, while the chlorophyll content in *LSU1OX-1* and *LSU1OX-3* plants was higher. The chlorophyll content in *lsu1-1* and *lsu1-2* mutants decreased by 36.9% and 27.5% respectively, while the chlorophyll content in LSU1OX-1 and LSU1OX-3 plants increased by 27.5% and 26.3% respectively (Fig. 4I). In short, these results confirmed that the loss of *LSU1* function led to a decrease in tolerance to Cd, while the overexpression of *LSU1* enhanced the tolerance to Cd.

We simultaneously treated Arabidopsis seedlings with 75 μM CdCl_2_ and 150 μM GSH. As for the PR length, the addition of GSH reduced the toxicity of Cd in Col-0, *lsu1-1*, *lsu1-2*, LSU1OX-1 and LSU1OX-3 seedlings, and the effects of GSH was most obvious in *lsu1-1* and *lsu1-2* mutants (Fig. 4G); the application of GSH promoted the FW of *lsu1-1* and *lsu1-2* plants (Fig. 4H), and there was no difference between Col-0 and *lsu1-1* and *lsu1-2* plants even under the condition of 75 μM Cd+ 150 μM GSH. A similar trend of chlorophyll content can be observed (Fig. 4I). These data suggest that the overexpression of *LSU1* enhances the tolerance of *Arabidopsis* to Cd via GSH at least in part. As documented, butylthionine sulfoxide (BSO) is glutathione synthesis inhibitor (Sheng et al., 2019). When treated with 75 μM Cd + 100 μM BSO, compared with Col-0, *lsu1-1* and *lsu1-2* mutants showed PR length reduction by 23.7% or 24.1% (Fig. 4G), fresh weight (FW) decreased by 21.3% or 21.2% (Fig. 4H), and chlorophyll content decreased by 28.6% or 26.2% (Fig. 4I). When compared with Col-0, no significant difference was observed between LSU1OX-1, LSU1OX-3, and Col-0 under the 75 μM Cd + 100 μM BSO (Fig. 4G H I). These results suggest that the role of LSU in Cd tolerance is dependent on biosynthesis of endogenous GSH.

To test the effects of *LSU1* on Cd accumulation, we measured the Cd content in roots and leaves of Col-0, *lsu1-1*, *lsu1-2*, LSU1OX-1 and LSU1OX-3 plants under Cd stress. Compared with Col-0, *lsu1-1*, *lsu1-2*, and mutants showed lower Cd content in roots and leaves, which decreased by 7.2% and 6.5% in roots and 9.5% (*P* <0.05) and 7.0% (*P <*0.05) in leaves, respectively. In contrast, LSU1OX-1 and LSU1OX-3 plants showed higher Cd content than Col-0, increasing by 17.9% (*P <*0.05) and 18.6% (*P <*0.05) in roots and 7.9% and 7.8% in leaves, respectively (Fig. S5A). We then identified the Cd transporter under Cd stress and observed that there was no difference between them (Fig. S5B). These results suggest that *LSU1* regulates Cd tolerance in *Arabidopsis thaliana* rather than by reducing Cd absorption or transport.

Next, we measured the levels of reduced glutathione (GSH), oxidized glutathione (GSSG), total glutathione, NPT, and phytochelatin (PC) in Col-0, *lsu1-1*, *lsu1-2*, LSU1OX-1 and LSU1OX-3 plants (Fig. S5 C-G). Cd treatment significantly increased GSH content in Arabidopsis. Under normal conditions, no significant differences in GSH content were observed between Col-0 and the *lsu1-1*, *lsu1-2*, LSU1OX-1, and LSU1OX-3 lines. However, after Cd treatment, GSH content decreased by 5.6% (*P <*0.05) and 6.6% (*P <*0.05) in *lsu1-1* and *lsu1-2*, respectively, while it increased by 13.3% (*P <*0.05) and 11.2% (*P <*0.05) in LSU1OX-1 and LSU1OX-3, respectively (Fig. S5 C).

In contrast, Cd treatment markedly reduced GSSG content in Arabidopsis. Under normal conditions, compared with Col-0, GSSG content decreased by 28.6% (*P <*0.05) and 17.4% (*P <*0.05) in *lsu1-1* and *lsu1-2*, respectively, and increased by 11.9% (*P <*0.05) and 13.6% (*P <*0.05) in LSU1OX-1 and LSU1OX-3, respectively. Following cadmium treatment, only LSU1OX-3 exhibited a 10.0% (*P <*0.05) increase in GSSG content relative to Col-0 (Fig. S5 D).

In addition, Cd has no effect on total glutathione in plants (Fig. S5E). However, in the absence of Cd, compared with Col-0, the total glutathione content of LSU1OX-1 and LSU1OX-3 increased by 7.74% and 8.61% respectively, while *lsu1-1* and *lsu1-2* decreased by 6.56% and 4.34% (*P* <0.05) respectively (Fig. S5E). Under Cd treatment, compared with Col-0, the contents of total glutathione in *lsu1-1* and *lsu1-2* mutants decreased by 6.89% and 6.24% (*P <*0.05) respectively, and the contents of LSU1OX-1 and LSU1OX-3 increased by 13.21% and 13.47% (*P <*0.05) respectively (Fig. S5E).

In the Cd-free medium, compared with Col-0, the NPT level in LSU1OX-1 and LSU1OX-3 plants increased by 4.7% (*P* <0.05) and 4.7% (*P* <0.05), respectively, while no difference was observed between *lsu1-1*, *lsu1-2*, and Col-0 (Fig. S5F). The content of NPT in Col-0, *lsu1-1*, *lsu1-2*, LSU1OX-1 and LSU1OX-3 plants increased under Cd stress. Compared with Col-0, the content of NPT in *lsu1-1* and *lsu1-2* decreased by 8.7% (*P* <0.05) and 7.6% (*P* <0.05) respectively, while the content in LSU1OX-1 and LSU1OX-3 plants increased by 20.7% (*P* <0.05) and 21.2% (*P* <0.05) respectively (Fig. S5F). In the absence of Cd, there was no significant difference in PC level among Col-0, *lsu1-1*, *lsu1-2*, LSU1OX-1 and LSU1OX-3 plants. However, under Cd stress, the PC contents of Col-0, *lsu1-1*, *lsu1-2*, LSU1OX-1 and LSU1OX-3 increased significantly. Compared with Col-0, the PC content of *lsu1-1* and *lsu1-2* plants decreased by 10.5% and 9.1%, while that of LSU1OX-1 and LSU1OX-3 plants increased by 25.2% and 25.9% (Fig. S5G). These results indicate that LSU1 positively regulates PC production under Cd stress.

As shown, WRKY45 binds to *LSU1* promoter and positively regulates its transcription (Fig. 3). We hypothesized that the phenotype of *wrky45* mutation under Cd stress could be rescued by overexpressing *LSU1*. To this end, we overexpressed LSU1 in *wrky45* mutant. qRT-PCR results verified that *wrky45*/LSU1OX-6 and *wrky45*/LSU1OX-7 plants can be used (Fig. 5A). Under control conditions, there was no significant difference in the growth of *wrky45*/LSU1OX-6 and *wrky45*/LSU1OX-7 plants compared with Col-0 and *wrky45* mutant plants (Fig. 5B). However, based on PR length (Fig. 5C), FW (Fig. 5D) and chlorophyll content (Fig. 5E) in 75 μM Cd^2+^, *wrky45*/LSU1OX-6 and *wrky45*/LSU1OX-7 plants were less sensitive to Cd than *wrky45* mutants and even more tolerant to Cd than Col-0 plants. Overall, these findings suggest that LSU1 acts as the downstream player of *WRKY45* genetically.

**Fig. 5.**
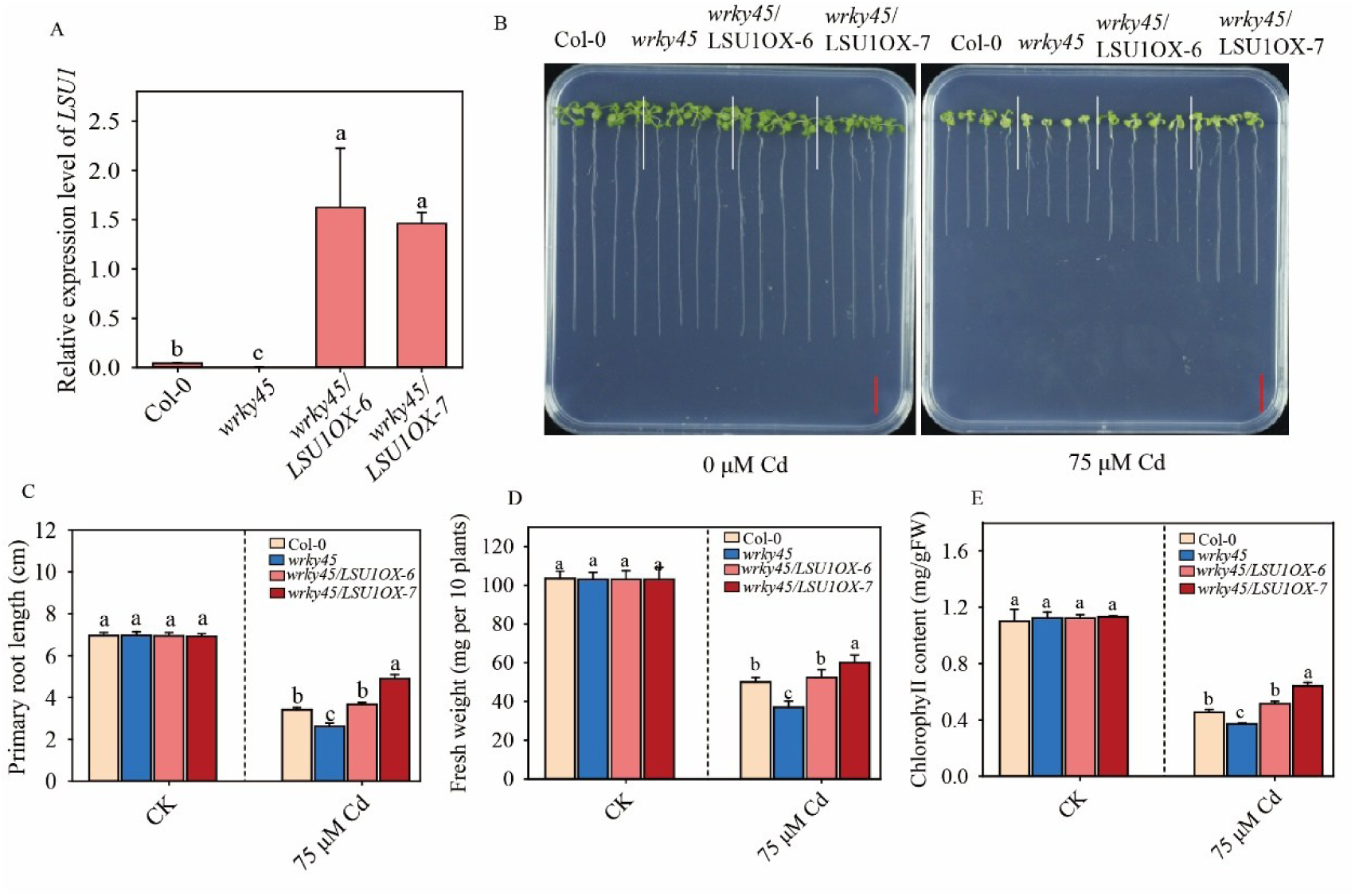
Overexpression of *LSU1* reduces the sensitivity of *wrky45* mutant to Cd. (A) Expression of *LSU1* of *wrky45*, wild type Col-0, *wrky45*/LSU1OX-6, and *wrky45*/ LSU1OX-7 lines. (B) Growth of Col-0, *wrky45*, *wrky45*/LSU1OX-6, and *wrky45*/ LSU1OX-7 lines under Cd stress. Two-day-old plants grown on 1/2 MS media were transferred to 1/2 MS media containing 0 or 75 μM CdCl_2_. Photographs were taken 10 days after treatment. Scale bar, 1 cm. Primary root length (C), fresh weight (D) and chlorophyll (E) of plants were measured. Three independent experiments were done with similar results, each with three biologicals repeats. Four plants per genotype from one plate were measured for each repeat. Data are presented as means ± SE (n = 4). Different letters indicate significant difference at *P <* 0.05 level (one-way ANOVA with Turkey’s test).

### 3.4 APS1 positively regulates cadmium tolerance through glutathione-dependent PC synthesis pathway

ATP sulfurylase (APS) is a key enzyme for plants to catalyze the transformation of inorganic SO_4_^2-^ into organic thioacyl glucoside sulfuric acid, and its catalytic action depends on ATP to provide energy (Hatzfeld et al.,2000). As documented, *LSU1* might interacts with ATP sulfurylase 1 (APS1) (Garcia-Molina et al. 2021). Hence, we speculate that there may be LSU1-APS1 interaction mechanism in Arabidopsis to regulate cadmium detoxification.

To test our hypothesis, we firstly studied the transcripts of *APS1* under Cd stress by qRT-PCR. Compared with the control, the expression of *APS1* increased significantly at 6 h and 24 h after Cd treatment. In Arabidopsis root, when treated with 6 h Cd, the transcription level of *APS1* was the highest, which was up-regulated by 1.4 times and 0.6 times respectively. In leaves treated with Cd for 24 hours, the transcription level of *APS1* was up-regulated by 1.2 and 0.5 times, respectively (Fig. 6AB). *APS2*, *APS3* and *APS4* are homologous genes of *APS1*. The transcription level of *APS2* in roots did not change after 6 h of Cd stress, but it was up-regulated after 24 h of Cd stress. The transcription level of *APS2* in leaves was up-regulated at 6 h and 24 h after Cd stress. The transcription level of *APS3* in roots and leaves was significantly up-regulated after 6 h and 24 h Cd stress. Similarly, the transcription level of *APS4* in the roots and leaves was significantly up-regulated after 6 h and 24 h Cd stress. These results revealed that the *APS1*, *APS2*, *APS3* and *APS4* respond to Cd stress (Fig. S6).

**Fig. 6.**
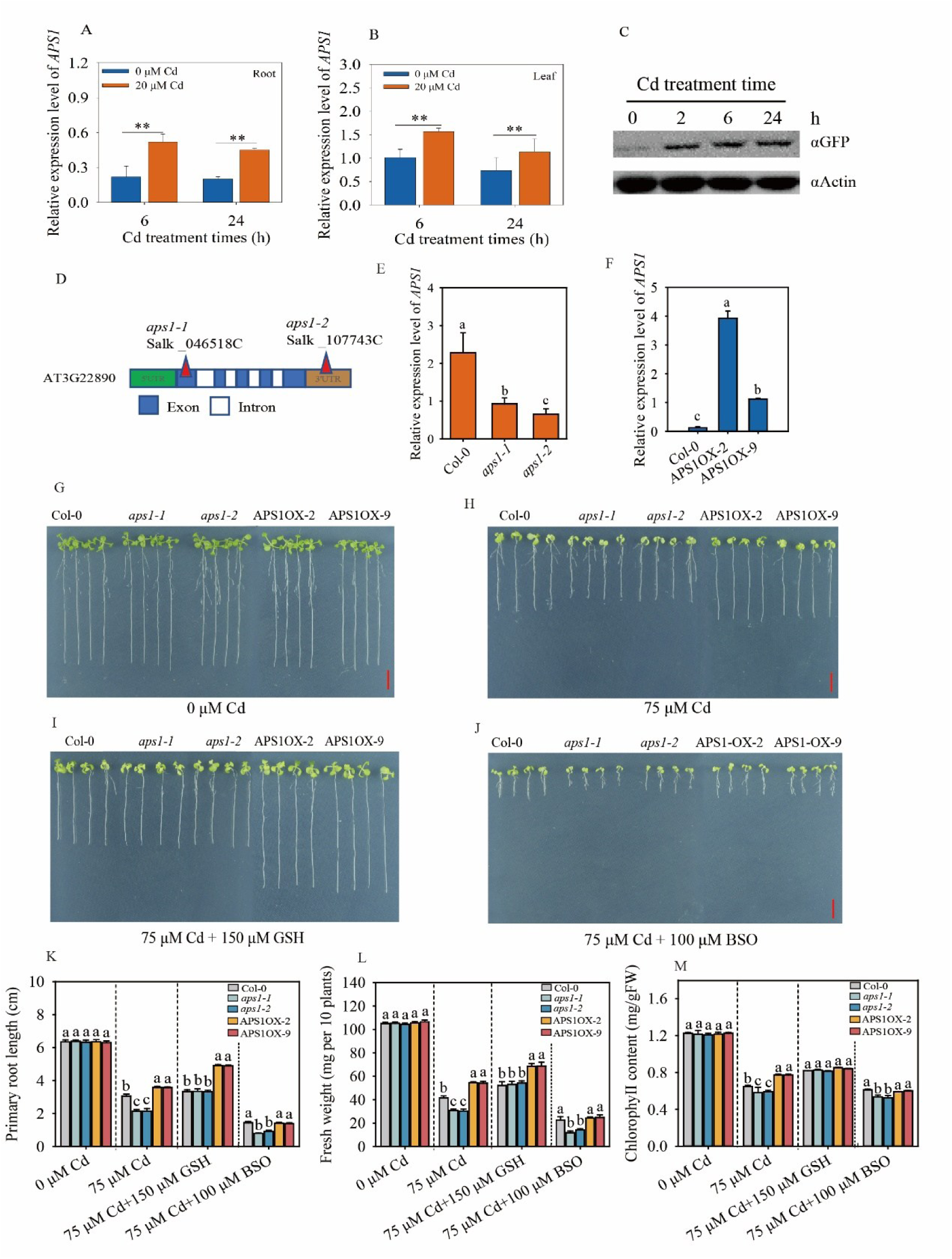
APS1 expression is promoted by Cd stress and positively regulates Cd tolerance in Arabidopsis. (A, B) The relative expression levels of *APS1* in leaves (A) and roots (B) of Arabidopsis were analyzed by qRT-PCR after 6 hours and 12 hours Cd stress, respectively. The relative expression level of *APS1* was normalized with *ACTIN2* (*AT3G18780*). (C) Western blotting analysis of APS1-GFP protein in ProAPS1::APS1-GFP plants under Cd stress. The plants were grown in nutrient solution for 2 weeks. After being treated with 20 μM CdCl_2_ for 0, 2, 6, and 24 hours, the total protein was extracted and analyzed using anti-GFP or anti-Actin antibodies. With Actin as the load control, the relative content of protein was quantitatively determined by ImageJ software. (D) Positionss of T-DNA insertion in *aps1-1* and *aps1-2* mutants. (E, F) Expression levels of *APS1* in the Col-0, *aps1-1*, *aps1-2* and overexpressing *APS1* APS1OX-2 and APS1OX-9 lines, respectively. Data represent means ± SD (n = 4) from four independent experiments. The asterisks indicate significant differences from the control (*P <* 0.01, Student’s *t*-test). (G-M) APS1-mediated Cd tolerance depended on endogenous glutathione biosynthesis. Arabidopsis was grown on 1/2 MS media containing 0 μM Cd (G), 75 μM Cd (H), 75 μM Cd+150 μM GSH (I), or 75 μM Cd + 100 μM BSO, respectively (J). The seedlings were grown on 1/2 MS media for 2 days, and then treated with 0 μM Cd, 75 μM Cd, 75 μM Cd+150 μM GSH, or 75 μM Cd + 100 μM BSO for 10 days, respectively. Scale bar, 1 cm. The primary root length (K), fresh weight (L) and chlorophyll content (M) of Col-0, *aps1-1*, *aps1-2*, APS1OX-2 and APS1OX-9 were measured. Three independent experiments were done with similar results. Four plants per genotype from one plate were measured for each repeat. Data present the means ± SE (n = 4). Different letters indicate significant difference at *P <* 0.05 level (one-way ANOVA with Turkey’s test).

In this study, ProAPS1:APS1-GFP transgenic line was used to study the protein expression level of APS1 by western blotting. The protein level of APS1 increases with the duration of Cd treatment. Within 2h of Cd treatment, the protein level of APS1 gradually increased, and it was found that the protein level of APS1 was 4.21, 4.82 and 5.21 times higher than that of the control at 2 h, 6 h and 24 h of Cd treatment, respectively. These results confirmed that Cd stress increased the protein level of APS1 (Fig. 6C).

To analyze the physiological function of *APS1* under Cd stress, documented two T-DNA insertion mutants of *APS1*, *aps1-1*(Salk_046518) and *aps1-2*(Salk_107743) were used in this study. The *aps1-1* mutant is produced by inserting T-DNA into the exon of *APS1*. The *aps1-2* mutant is produced by inserting T-DNA into the 3’ untranslated region of *APS1* (Fig. 6D). The expression levels of APS1 in *aps1-1* and *aps1-2* mutants were 40.2% and 28.3% of Col-0, respectively (Fig. 6E). In addition, the overexpression materials *APS1*OX-1, *APS1*OX-3, *APS1*OX-4, *APS1*OX-7 were obtained by transgenic technology. The transcription level of *APS1*OX-2 and *APS1*OX-9 materials is 30.4 and 11.1 times higher than that of Col-0, respectively (Fig. 6F).

Compared with Col-0, no significant growth difference was observed in the strains of *aps1-1*, *aps1-2*, *APS1*OX-2 and *APS1*OX-9 grown in 1/2MS medium (Fig. 6G). However, overexpressing *APS1* showed higher Cd tolerance than Col-0, while *aps1* mutant was more sensitive to Cd (Fig. 6H). When treated with 75 μM CdCl_2_, the PR length of *aps1-1* and *aps1-2* mutants was significantly reduced by 30.1% and 31.2% compared with Col-0. On the contrary, the PR length of *APS1*OX-2 and *APS1*OX-9 increased by 17.2% and 16.9% respectively (Fig. 6K). The FW of *aps1-1* and *aps1-2* decreased by 25.8% and 27.1% respectively, while the FW of *APS1*OX-2 and *APS1*OX-9 plants increased by 25.3% and 26.1% respectively (Fig. 6L). When treated with 75 μM CdCl_2_, the chlorophyll content of *aps1-1* and *aps1-2* was lower than that of Col-0. However, the chlorophyll content of *APS1*OX-2 and *APS1*OX-9 is higher. The chlorophyll content in *aps1-1* and *aps1-2* decreased by 10.1% and 8.9% respectively, while the chlorophyll content in *APS1*OX-2 and *APS1*OX-9 plants increased by 18.1% and 19.2% respectively (Fig. 6M). These results address that *APS1* positively regulates Cd tolerance.

*APS1* may affect GSH and PC synthesis by participating in the assimilation of SO_4_^2-^. The Arabidopsis seedlings were treated simultaneously with 75 μM Cd^2+^ and 150 μM GSH. Addition of GSH could mitigate the symptoms of Cd toxicity in Col-0, *aps1-1*, *aps1-2*, *APS1*OX-1 and *APS1*OX-3 seedlings, and the effect of GSH was more obvious in the *aps1-1* and *aps1-2* mutants. The application of GSH increased the fresh FW of *aps1-1* and *aps1-2* (Fig. 6L). Under the condition of 75 μM Cd^2+^ plus 150 μM GSH, there was no difference between Col-0, *aps1-1* and *aps1-2*. A similar trend can also be observed in chlorophyll content (Fig. 6M). These findings suggest that *APS1* regulates Cd tolerance via affecting biosynthesis of endogenous GSH.

When treated with 75 μM Cd^2+^ plus 100 μM BSO, the PR length of the *aps1-1* and *aps1-2* mutants was reduced by 44.8% and 37.0% (Fig. 6K), the FW was reduced by 48.1% and 37.3% (Fig. 6L), and the chlorophyll content was reduced by 12.1% or 13.8% (Fig. 6M) compared with Col-0. Under the same conditions, no significant differences were observed between *APS1*OX-2, *APS1*OX-9 and Col-0 (Fig. 6G-M). These results show that overexpression of *APS1* enhances the tolerance of Arabidopsis to Cd through GSH synthesis pathway.

Compared with Col-0, the Cd content in the roots of *aps1-1* and *aps1-2* mutants decreased by 6.4% and 8.6% respectively, and 9.1% and 8.2% respectively in leaves, respectively; in contrast, the Cd content in APS1OX-2 and APS1OX-9 roots increased by 11.7% and 13.7%, respectively, and 7.0% and 8.3% respectively in leaves (Fig. S6A). In addition, there was no difference in the Cd transport coefficients of Col-0, *aps1-1*, *aps1-2*, APS1OX-2 and APS1OX-9 under Cd treatment (Fig. S6B). This indicates that APS1 does not affect the redistribution of Cd between roots and leaves.

Under normal conditions, no significant differences in GSH content were detected between Col-0 and the *aps1-1*, *aps1-2*, APS1OX-2, and APS1OX-9 lines. Following Cd treatment, however, GSH content decreased by 20.9% (*P <*0.05) and 21.5% (*P <*0.05) in *aps1-1* and *aps1-2*, respectively, whereas it increased by 6.1% (*P <*0.05) and 7.5% (*P <*0.05) in APS1OX-2 and APS1OX-9, respectively (Fig. S6C). In contrast, Cd exposure markedly reduced GSSG levels in Arabidopsis. Under control conditions, GSSG content was 24.9% (*P <*0.05) and 25.5% (*P <*0.05) lower in *aps1-1* and *aps1-2*, respectively, relative to Col-0 (Fig. S6D).

In the absence of Cd, total glutathione content was elevated by 14.7% and 15.3% in APS1OX-2 and APS1OX-9, respectively, while it declined by 8.2% and 8.5% in *aps1-1* and *aps1-2*, respectively (Fig. S6E). Under Cd treatment, total glutathione content decreased by 8.2% and 9.4% (*P <*0.05) in the *aps1-1* and *aps1-2* mutants, respectively, and increased by 14.6% and 15.3% in APS1OX-2 and APS1OX-9, respectively (Fig. S6E).

Under normal conditions, the NPT content of APS1OX-2 and APS1OX-9 increased by 10.4% and 8.9%, respectively, while the NPT content of *aps1-1* and *aps1-2* decreased by 6.7% and 9.9%, respectively (Fig. S6F). Compared with Col-0 under Cd treatment, the NPT content in *aps1-1* and *aps1-2* decreased by 16.4% and 16.2% respectively, while the NPT content in APS1OX-2 and APS1OX-9 plants increased by 18.6 and 15.0% respectively (Fig. S6F).

In addition, under the absence of Cd, there was no significant difference in PC levels of Col-0, *aps1-1*, *aps1-2*, APS1OX-2 and APS1OX-9. However, under Cd stress, the PC content of Col-0, *aps1-1*, *aps1-2*, APS1OX-2 and APS1OX-9 increased significantly. Compared with Col-0 under Cd stress, the PC content of *aps1-1* and *aps1-2* was reduced by 22.7% and 20.4%, while the PC content of *APS1*OX-2 and *APS1*OX-9 plants increased by 27.1% and 19.5% (Fig. S6G). These results suggest that *APS1* positively PC synthesis under Cd stress.

### 3.5 LSU1 interacts with APS1 to enhance ATP sulfurylase activity

To verify the interaction between APS1 and LSU1, we expressed and purified LSU1-His and GST-APS proteins, respectively. The physical interaction between APS1 and LSU1 was further proved through in *vitro* pull-down experiments (Fig. 7A). Compared with the co-incubation of control GST protein and LSU-His protein, after co-incubation of GST-APS1 protein and LSU-His protein, the obvious LSU-His western blot can be detected in the eluent by binding to GST magnetic beads.

**Fig. 7.**
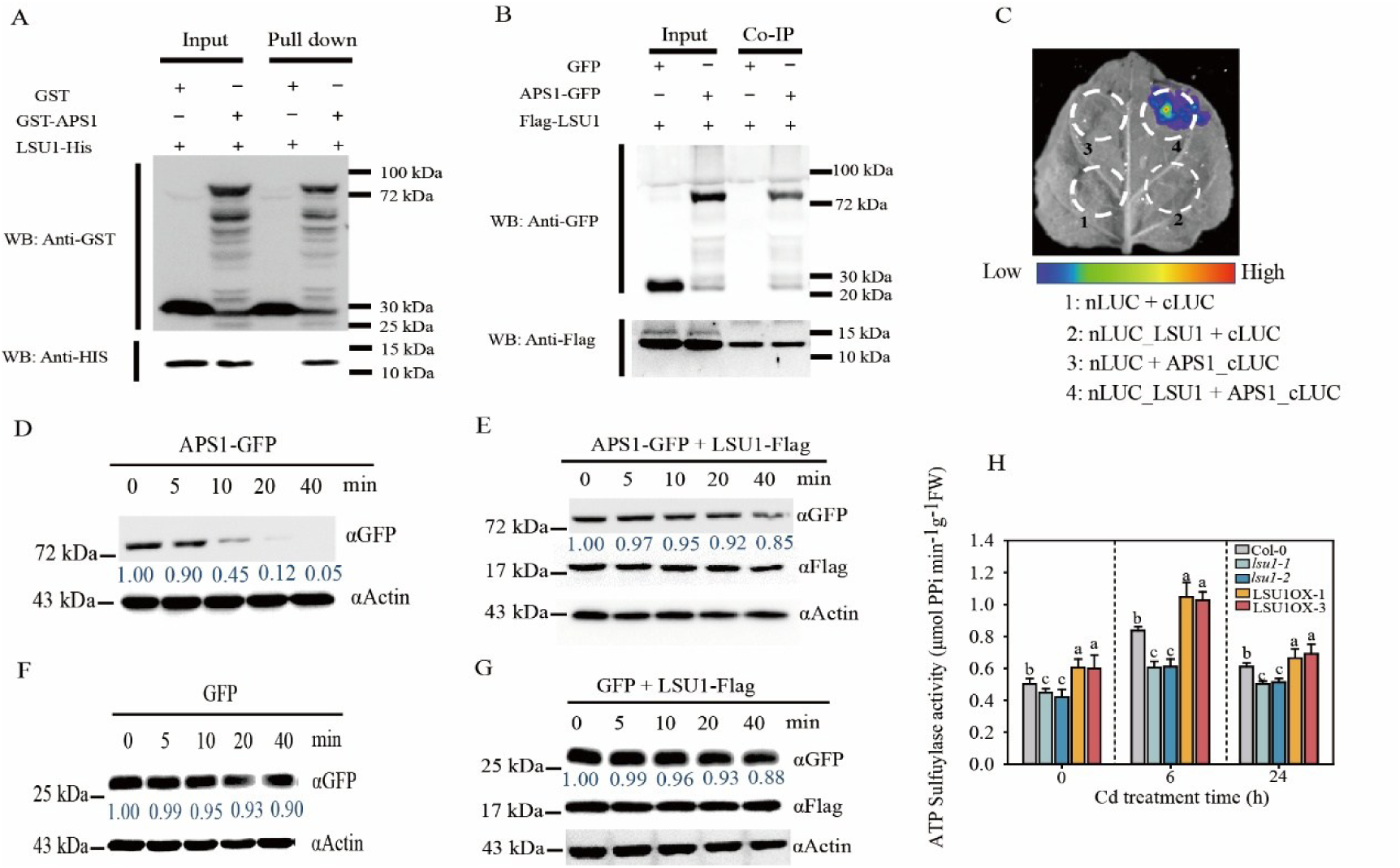
LSU1 physically interacts with APS1 both *in vitro* and *in vivo*, slows down the degradation rate of the APS1 protein, and enhances ATP sulfurylase activity. (A) Pull-down assay for the interaction between LSU1 and APS1. GST, APS1-GST and LSU-His were separately expressed and purified in *Escherichia coli* and then used for GST pull-down assays. Anti-GST and anti-His antibodies were used to detect GST, APS1-GST, and LSU1-His, respectively. (B) Co-IP assay for interaction between LSU1 and APS1 in Arabidopsis. APS1-GFP, LSU1-flag and GFP empty vector were transfected into Arabidopsis, and protein extracts were immunoprecipitated (IP) using an anti-Flag antibody and were detected with anti-Flag (LSU1-Flag) and anti-GFP (APS1-GFP and GFP) antibodies. Protein input is shown by immunoblot analysis of protein extracts before immunoprecipitation and antibodies against the respective tags. Molecular mass markers are shown on the right. (C)Split-LUC assay for interaction between LSU1 and APS1 in *N. benthamiana* leaves. Full-length LSU1 protein was fused to the N terminus of firefly LUC (LSU1-nLUC), and full-length APS1 was fused to the C terminus (APS1-cLUC). Co-infiltration of nLUC and cLUC, LSU1-nLuc and cLUC, or nLUC and APS1-cLUC were used as negative control treatments. (D-G) LSU1 attenuates the degradation rate of APS1 protein in Arabidopsis. The plants were transfected with APS1-GFP, LSU1-Flag, and GFP empty vectors, and total proteins were extracted. The plant proteins containing APS1-GFP (D), APS1-GFP+LSU1-Flag (F), GFP (D), and GFP+LSU1-Flag (B) were incubated at 22℃. The samples after incubation were examined using Western blotting. Anti-GFP and Anti-Flag antibodies were used to detect GFP, APS1-GFP, and LSU1-Flag, respectively. (H) LSU1 increases the ATP sulfurylases activity of APS1. Col-0, *lsu1-1, lsu1-2*, LSU1-OX-1 and LSU1-OX-3 seedlings were grown in nutrient solution for 2 weeks, and after treated with 0 or 20 µM CdCl2 for 24 h, then the ATP sulfurylases activities were quantified. Data present the means ± SE (n = 4). Different letters indicate significant difference at *P <* 0.05 level (one-way ANOVA with Turkey’s test).

We also obtained stable trans genetic lines of 35S::APS1-GFP, 35S::GFP and 35S::LSU1-Flag. LSU1-Flag, GFP and APS1-GFP proteins were extracted from Arabidopsis seedlings, respectively. Compared with the co-incubation of control LSU1-Flag and GFP protein, after co-incubation with APS1-GFP protein, a significant APS1-GFP western blot can be detected in the eluent by binding to Flag magnetic beads. Co-IP analysis confirmed that LSU1 interacts with APS1 *in vivo* (Fig. 7B). We also verified the interaction of AtLSU1 with AtAPS1 by split luciferase (Split-LUC) experiment (Fig. 7C). Compared with the negative controls of co-converted *Nicotiana benthamiana* with nLUC, LSU1-nLUC with cLUC and nLUC with APS1-cLUC, obvious LUC fluorescence could be observed.

To further understand the biochemical importance of the interaction between LSU1 and APS1. Total APS1-GFP protein in transgenic plants was extracted and incubated at 22°C, and GFP antibodies were used for detection. We found that after 40 minutes of incubation, APS1-GFP was degraded over time, 0.05 times that of the initial loading protein (Fig. 7D). Moreover, after the purified LSU1-Flag was added to the total APS1-GFP protein, the degradation rate of APS1-GFP protein decreased. After 40 minutes of incubation, the APS1-GFP protein was 0.85 times that of the initial loading protein, which was much higher than the control without LSU1-Flag (Fig. 7E). In order to exclude the experimental results of the degradation of GFP protein, a GFP protein degradation experiment was set up separately and found that the degradation rate of GFP protein was much lower than that of APS1-GFP protein, which was 0.90 times that of the initial loading protein after 40 minutes incubation (Fig. 7F); in addition, after the addition of LSU1-Flag did not slow down the GFP protein turnover (Fig. 7G). To sum up, LSU1 interacts with APS1 to slow down APS1 degradation, thus increasing APS1 stability.

We thus detected the ATP sulfurylase activity of Col-0, *lsu1-1*, *lsu1-2*, *LSU1*OX-1 and *LSU1*OX-3 at Cd treatment of 0 h, 6 h, and 24 h. Cd stress increased ATP sulfurylase activity, and increased first and then decreased within 24 h of Cd treatment, and peaked after 6 h of Cd treatment. *LSU1* can increase ATP sulfurylase activity. Under 0 h Cd treatment, the ATP sulfurylase activity levels in *LSU1*OX-1 and *LSU1*OX-3 plants increased by 20.4% and 19.4%, respectively, while the ATP sulfurylase activity levels in *lsu1-1* and *lsu1-2* decreased by 10.8% and 16.1% respectively. After 6 h of Cd treatment, compared with Col-0, the ATP sulfurylase activities in *lsu1-1*, *lsu1-2* and 27.1% were reduced (*P <*0.05), while the ATP sulfurylase activities of *LSU1*OX-1 and *LSU1*OX-3 increased by 25.2% and 22.6% respectively (Fig. 7H). In addition, after 24 h of Cd treatment, compared with Col-0, the ATP sulfurylase activity in *lsu1-1* and *lsu1-2* was reduced by 17.7% and 15.9% (*P<*0.05), respectively, and the content of *LSU1*OX-1 and *LSU1*OX-3 in plants increased by 8.8% and 13.3% (Fig. 7H), respectively. These results show that LSU1 increases ATP sulfurylase enzyme activity at physiological level.

To verify whether *LSU1* interacts with *APS1* genetically, we introduced *LSU1*OX-3 into the *aps1-1* mutant through crossing. Overexpressing *LSU1* lines were screened in the *aps1-1* background by qRT-PCR (Fig. 8A). In Cd-free 1/2 MS media, no significant differences were found in the growth of *aps1-1*/*LSU1*OX-3 plants compared with the Col-0, *APS1*OX-9 and *aps1-1* mutant plants (Fig. 8B). However, in 1/2 MS media containing 75 μM Cd^2+^, *aps1-1* and *aps1-1*/*LSU1*OX-3 were sensitive to Cd compared with the Col-0 and *LSU1*OX-3 mutants, and overexpression of *LSU1* in *aps1-1* mutant did not restore its Cd-sensitive phenotype (Fig. 8C D E). Therefore, we conclude that *LSU1* is dependent on *APS1* to regulate Cd tolerance.

**Fig. 8.**
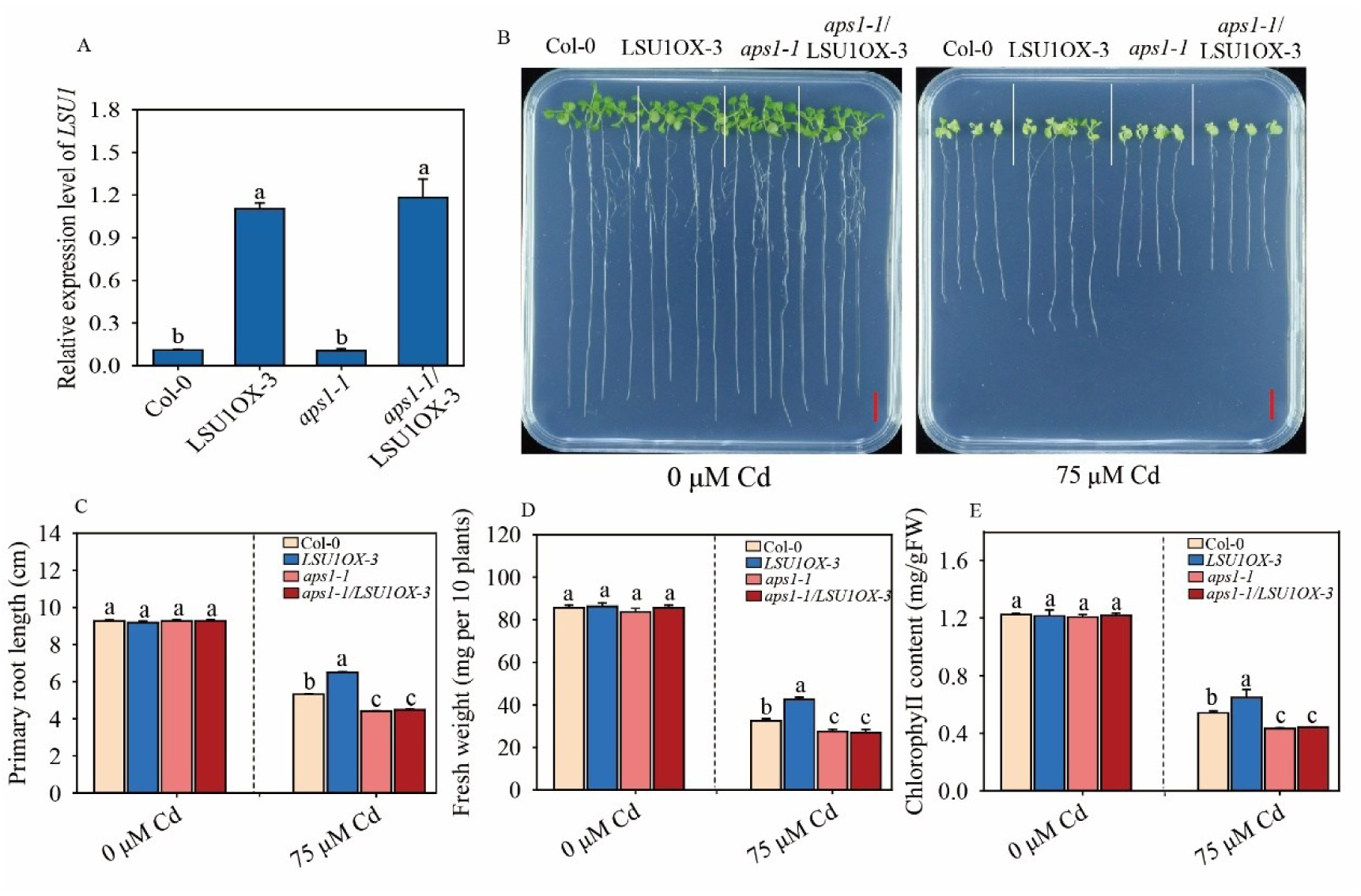
Overexpression of *LSU1* did not reduce the Cd sensitivity of *aps1-1* mutants. (B) Growth of Col-0, LSU1OX-3, *aps1-1*, and *aps1-1*/ LSU1OX-3 lines under Cd stress. Two-day-old plants grown on 1/2 MS media were transferred to 1/2 MS media containing 0 or 75 μM CdCl_2_. Photographs were taken 10 days after the transfer. Scale bar, 1 cm. Primary root length (C), fresh weight (D) and chlorophyll (E) of plants were measured. Three independent experiments were done with similar results, each with three biologicals repeats. Four plants per genotype from one plate were measured for each repeat. Data are presented as means ± SE (n = 4). Different letters indicate significant difference at *P <* 0.05 level (one-way ANOVA with Turkey’s test). (E)

**Fig. 9.**
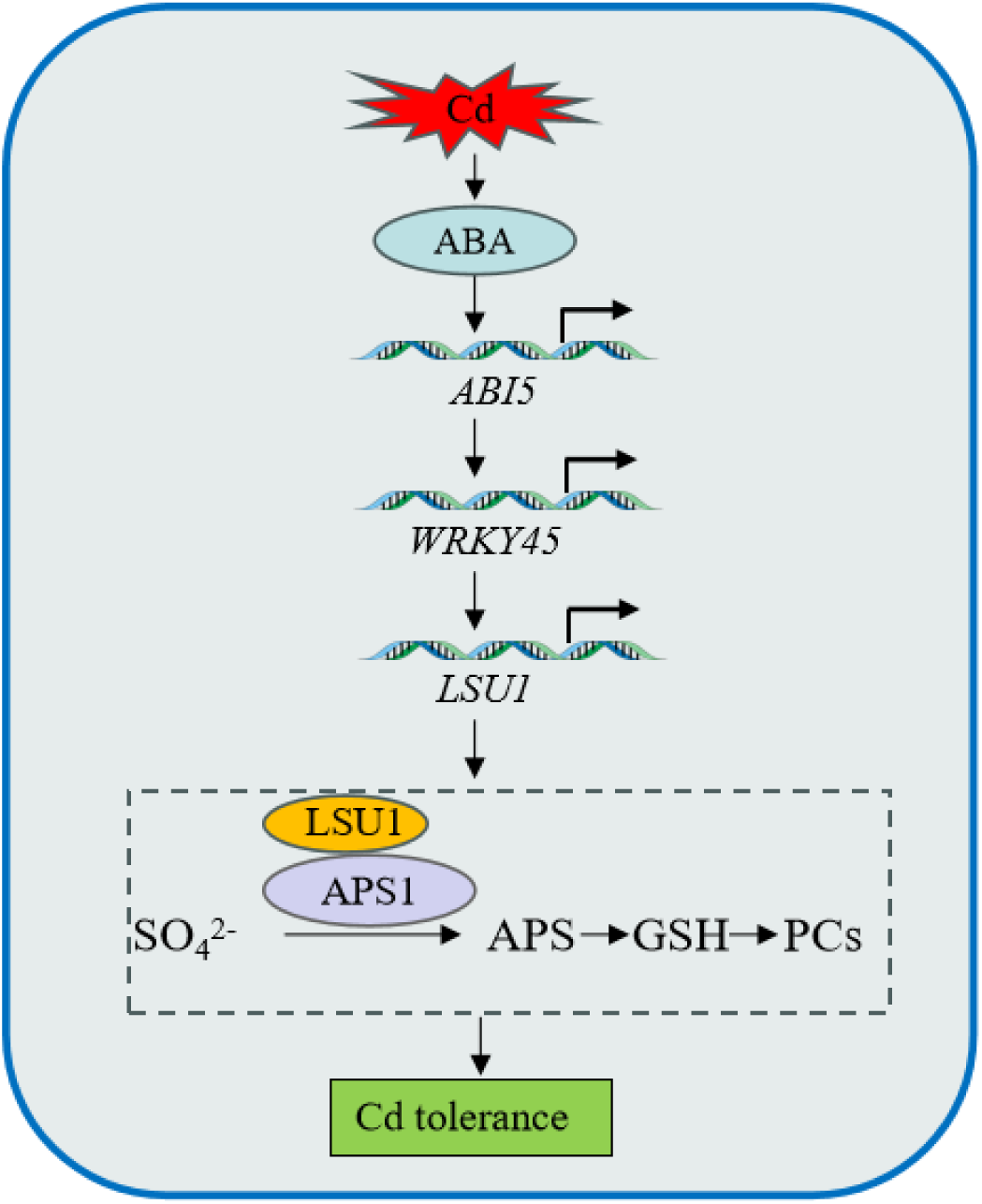
The working model of ABI5-WRKY45-LSU1 in regulating Cd tolerance. Cadmium stress triggers ABA accumulation and subsequent *ABI5* expression. ABI5 directly binds to the ABRE motif in the *WRKY45* promoter and activates its transcription. *WRKY45* targets *LSU1*, whose overexpression enhances Cd tolerance whereas knockout confers sensitivity. LSU1 stabilizes APS1 to elevate ATP sulfurylase activity, thereby promoting GSH and PC accumulation.

## 4. Discussion

Cd stress increases the levels of endogenous ABA in rice, potato, kale rape and wheat (Hsu et al., 2010; Stro et al., 2013). In addition, ABA to alleviates Cd toxicity Arabidopsis, mung beans (*Vigna Radiata L.*) and poplars (*Pillius Euphatica*) (Han et al., 2016; Pan et al., 2020; He et al., 2021). Arabidopsis ABA synthesis-deficient mutants *aba3*, *aba-4*, *nced3*, and ABA insensitive mutants *abi2-1*, *abi3-1* are sensitive to Cd stress and accumulate more Cd (Sharma et al., 2002). In this study, we revealed that ABI5-WRKY45-LSU1 axis confers Cd tolerance.

This study revealed that Cd stress upregulates the expression of ABA synthetic genes *NCED3*, *ABA2*, and *AAO3* in Arabidopsis, and inhibits the expression of ABA degraded genes *CYP707A1* and *CYP702A2,* respectively (Fig. S1). Cadmium treatment can increase endogenous ABA content in Arabidopsis (Fig. S1A). In line with previous studies, application of 0.5 μM ABA can effectively alleviate Cd toxicity (Fig. S1). As reported, low concentrations of ABA can increase the content of polysaccharides and uronic acid in root cell walls and increase root Cd adsorption, increase plant Cd tolerance (Yu et al., 2023), inhibit *IRT1* expression in roots (Fan et al., 2014), and increase the expression of *PDF2.6*, *PDR8* and *AIT1* related to Cd chelation and efflux, reducing Cd accumulation and Cd toxic symptoms (Meng et al., 2022).

During Cd stress, ABA levels increase, which subsequently induces binding of PP2Cs to ABA receptors PYR/PYL/RCAR, thereby relieving the inhibition of SnRK2; SnRK2 can phosphorylate and activate downstream effector molecules such as crucial transcription factors *ZAT6* and *ABI5* (Ng et al., 2014). *ZAT6* forwardly regulates the expression of the target gene *GSH1* to increase GSH and PC synthesis (Chen et al., 2016). As shown, ABA or Cd treatment can increase transcription levels of *ABI5* and *WRKY45* (Fig. 1B and S2). Moreover, the *abi5-8* mutant is sensitive to Cd stress (Fig.1C-H). These results thus verified that ABI5 positively regulates Cd tolerance.

ABA regulates the expression of about 10% of editing protein genes in Arabidopsis, and the expression of these genes is mainly regulated by two different families of *bZIP* transcription factor *AREBs*/*ABFs* (Fujita et al., 2011). SnRK2s kinase phosphorylates and activates *ABF*s/*AREB*s, including *ABI3*, *ABI4*, and *ABI5* (Yang et al., 2018). *ABI5* inhibits the expression of *MYB49* and the downstream gene *bHLH38/39/100/101* by interacting with MYB49, thereby negatively regulating the transcription of *IRT1* and reducing Cd absorption; consistently the overexpression of *ABI5* increases the Cd tolerance of Arabidopsis (Zhang et al., 2019). *ABI5* forwardly regulates *PHO1* transcription involved in ABA-mediated seed germination and early seedling development (Huang et al., 2017); ABI5 binds ABRE cis-acting elements of the *WRKY42* promoter region to regulate *WRKY42* expression in response to low phosphorus stress (Lei et al., 2022). Here, we found that ABI5 directly binds to the ABRE element of the *WRKY45* promoter and forwardly regulates *WRKY45* transcription (Fig. 1). *ABI5* and *WRKY45* play a positive regulatory role in phosphorus nutrient absorption (Wang et al., 2014; Chen et al., 2017; Barros et al., 2022) and plant leaf aging (Liebsch et al., 2016). Therefore, it needs to be verified that the ABI5-WRKY45 module may play a role in regulating Arabidopsis’s phosphorus absorption and leaf aging.

The role of *MdABI5* in apple in promoting leaf aging can be enhanced through interactions with *MdWRKY44* and *MdbZIP44* (An et al., 2021), which shows that WRKY interaction with *ABI5* at the protein level is also a new regulatory mechanism. Accordingly, we can not rule out the possibility of ABI5 interacts with WRKY45 at protein level. In addition, whether the ABI5-WRKY45 module regulates Cd tolerance is conservative in rice, wheat, soybean and other crops still needs to be studied.

Under Cd stress, overexpression of *WRKY45* promotes the expression of PC synthase genes *PCS1* and *PCS2* (Li et al., 2023). Here, we uncover LSU1 is the novel target of WRKY45. The GO enrichment analysis of the differential genes between *WRKY45*OX-15, *wrky45* and Col-0 was found, and some related genes with sulfur assimilation were found (Fig. 3). The transcripts of *LSU1* and *LSU2* are significantly induced by low sulfur (S), low iron and copper stress (Garcia-Molina et al., 2017). *LSU1* and *LSU2* promote the degradation of glucosinolates by inhibiting the biosynthesis of glucosinolates, thereby limiting sulfur consumption, promoting the production of GSH and PC, and increasing the Cd tolerance of Arabidopsis (Li et al., 2023).

Cd stress induced transcription of *LSU1*, *LSU2* and *LSU3*, but its effects on *LSU4* was not obvious (Fig. S4). This study demonstrates that overexpressing *LSU1* can increase the tolerance of Arabidopsis to Cd, while mutants *lsu1-1* and *lsu1-2* are sensitive to Cd (Fig. 4). In addition, BSO treatment can reduce Cd tolerance in overexpressing *LSU1* strains (Fig. 4), and GSH treatment can restore Cd sensitivity of *lsu1* mutant (Fig. 4). This indicates that roles of LSU1 in Cd tolerance is dependent on GSH biosynthesis. Moreover, the contents of NPT, GSH, and PC content under Cd stress are significantly increased in overexpressing *LSU1* lines, but not in *lsu1* mutant (Fig. 4). Of note, overexpression of *LSU1* significantly increased the Cd content in Arabidopsis roots and leaves under Cd stress, while the opposite effect was observed in the *lsu1* mutant (Fig. S5). This also shows that LSU1-mediated Cd tolerance does not depend on the decrease of Cd absorption. LSU1 promotes GSH and PC synthesis, and main root elongation indirectly leads to an increase in Cd absorption.

The *LSU1* promoter contains bZIP transcription factor binding element (G-box), auxin, ethylene and jasmonic acid reaction elements, SQUAMOSA promoter binding element (SBP), and WRKY binding element (W-box) (Sirko et al., 2015). *EIN3* is an important member of ethylene signal transduction. *EIN3* targets negatively regulating the transcription of *XTH33* and *LSU1*, inhibiting the growth of Arabidopsis main root under Cd stress (Kong et al., 2018). Dual-LUC and EMSA data verified that *WRKY45* positively regulates *LSU1* transcription by directly binding to the W-box1 element of the *LSU1* promoter (Fig. 4). Moreover, overexpression of *LSU1* in *wrky45* mutant restores sensitivity to Cd stress (Fig. 5). This suggests that *LSU1* functions downstream of *WRKY45*.

*LSU1* encodes a short peptide of 94 amino acids and is localized to multiple organelles, including cytoplasm and microsomes, chloroplasts (Antoni et al., 2017). LSU1 has not enzyme activity or transcription regulatory functions (Rodrigues et al., 2019). Iron-dependent superoxide dismutase 2 (*FSD2*) physically interacts with *LSU1* and *LSU2 in vitro* and *in vivo*, which increases the enzymatic activity of *FSD2*, resulting in hydrogen peroxide (Garcia-Molina et al., 2017). IP-MS analysis show that APS1, GRF8, RAF2, and CAT2 are interacting proteins of LSU1 (Niemiro et al., 2020). Given that APS1 is the crucial enzyme involved in S assimilation, we hypothesized that LSU1 interacts with APS1 to affect S assimilation, thus boosting Cd tolerance. We firstly verified the interaction of LSU1 with APS1 based on Co-IP and split-LUC experiments (Fig. 7). Secondly, we revealed that overexpression of *LSU1* improves ATP sulfurylase activity, and mutations in *LSU1* gene cause a decrease in ATP sulfurylase activity (Fig. 7). At biochemical level, interaction of LSU1 with APS increases the stability of APS1, but the concrete mechanisms remain elusive.

ATP sulfurylase is a key enzyme that catalyzes the conversion of SO_4_^2^- in the inorganic state to sulfate in the organic state, and relies on ATP to provide energy (Vierstra, 2003). Overexpression of APS in soybeans can significantly increase the Cys content of seeds (Kim et al., 2020), which is an important precursor for plant synthesis of GSH and PCs (Kim et al., 2019). There are four APS genes in the Arabidopsis genome, namely *APS1*, *APS2*, *APS3*, and *APS4* (Bohrer et al., 2015). The transcription level of APS genes increases rapidly under Cd stress (Niemiro et al., 2020). This study showed that after 6 h of Cd treatment, the transcription level of *APS1* increased rapidly; and the protein level of APS1 continued to increase within 24 h of Cd stress (Fig. 6). Considering the possibility of post-translational modification of APS1 protein, it is necessary to explore kinases and E3 ligases responsible for their phosphorylation, ubiquitination and other modifications under Cd stress. The APS gene expression and ATP sulfurylase activity were higher in Cd superaccum plants such as Southeast Sedum and Centipede. It is worth noting that mutations in the APS1 gene in Arabidopsis increase the sensitivity of Arabidopsis to Cd, while overexpressing APS1 enhances Cd tolerance (Fig. 6). Therefore, APS1 positively regulates Cd tolerance of Arabidopsis. BSO treatment can reduce Cd tolerance in overexpressed APS1 strains (Fig. 6); GSH treatment can restore Cd sensitivity of aps1 mutant (Fig. 6). Overexpression of *APS1* significantly increased the NPT, GSH, and PC content under Cd stress, while the mutant APS1 was the opposite (Fig. S6). Therefore, it can be clarified that Arabidopsis *APS1* increases the synthesis of NBT, GSH, and PC by regulating the assimilation process of SO4^2-^, and regulates the detoxification of Cd toxicity in plants.

*APS2*, *APS3* and *APS4* are highly homologous to APS1 (Bohrer et al., 2015). The transcript levels of *APS2*, *APS3* and *APS4* are upregulated by Cd stress (Fig. S7). As reported, miR395 is a low-S-induced microRNA, and Arabidopsis APS1 is the target gene of miR395 (Bohrer et al., 2015). Whether *ABI5* and *WRKY45* regulate *miR395* expression in Cd stress is unclear. Under Cd conditions, how *miR395* regulates *APS1* at temporal and spatial pattern are worthy of future study.

IP-MS LSU1 interacts with APS1 (Niemiro et al., 2020), but its biological significance is unclear. This study proves that both APS1 and LSU1 participate in the regulation of Cd tolerance in plants through the GSH-PC synthesis pathway, so in-depth analysis of the mechanism of the interaction between LSU1 and APS1 is of great significance in the Cd tolerance of Arabidopsis. Studies have shown that Arabidopsis LSU2, LSU3 and LSU4 can form dimers with their own proteins respectively, and different combinations of heterodimers can be formed between LSU1-LSU4. Y2H shows that the binding between LSU1-LSU2 is relatively strong, while the binding between LSU4-LSU1 is relatively weak. The three-dimensional structure simulates the binding of LSU1-LSU1 homodimer, revealing that cysteine position 54 (C54) is more critical for dimer formation, and the Y2H experiment also proves this. BiFC experiments reveal that APS1 interacts with LSU1, LSU2, LSU3, and LSU4, and GRF8 interacts with LSU1, LSU2, LSU3, and LSU4 interacts, RAF2 interacts with LSU1, LSU2, and LSU4; Y2H confirms that CAT2 and NBR1 also interact with LSU1, LSU2, LSU3, and LSU4. Amino acid mutation analysis revealed that the L60A mutation of LSU1 significantly reduced the interaction between LSU1 and NBR, while the C54A, C54E, C54R, and L60A mutations affect the interaction between LSU1 and CAT and RAF2, but the C54E mutation with the greatest effect was the C54E mutation (Niemiro et al., 2020). These results provide cues for further studies on the interaction sites between LSU1 and APS1. Although we revealed that LSU1 interacts with APS1, it is not clear whether LSU1 interacts with APS2, APS3, and APS4. In addition, it needs to be deciphered that the key amino acid sequence determines the interaction between LSU1 and APS1 and where the interaction taken place at subcellular level.

Plant proteins are degraded by 26S ubiquitination and lysosomes in cell (Vierstra, 2003; Lazaro et al., 2012). Presently, how APS1 is degraded is unclear. Here, we revealed that adding LSU1 protein can slow down the degradation rate of APS1 protein *in vitro* (Fig.7). Thus, LSU1 may have changed the conformation of the APS1 protein through interaction with APS1, increasing APS1 stability, thereby boosting its ATP sulfurylase activity.

Therefore, it is very important to find the interaction sites between LSU1 and APS1 protein at the protein level and understand the specific mechanisms by which LSU1 slows down the degradation of APS1 protein. In addition, overexpression of LSU1 in the *aps1* mutant did not alter the sensitivity of *aps1* to Cd stress (Fig.8), indicating that the *LSU1* regulates Cd tolerance is APS1-dependent.

## 5. Conclusion

Cadmium stress triggers ABA accumulation and subsequent *ABI5* expression. ABI5 directly binds to the ABRE motif in the *WRKY45* promoter and activates its transcription. *WRKY45* targets *LSU1*, whose overexpression enhances Cd tolerance whereas knockout confers sensitivity. LSU1 stabilizes APS1 to elevate ATP sulfurylase activity, thereby promoting NPT, GSH and PC accumulation. Collectively, the ABI5–WRKY45–LSU1 module confers Cd tolerance in Arabidopsis by regulating sulfur metabolism and downstream thiol compound biosynthesis.

## CRediT authorship

Fangjian Li: Conceptualization, Methodology, Validation, Formal analysis, Funding acquisition, Project administration, Investigation, writing original draft, Writing review & editing. Xinni Zheng: Validation. Yanan Zhang: Validation. Jianjian Chen: Validation. Guihua Lv: Review. Jinxiang Wang: Supervision, Conceptualization, Funding acquisition, Project administration, Resources, Writing – review & editing.

## Data Availability

Data will be made available on request.

## Acknowledgements

This research was funded by the Zhejiang Provincial Natural Science Foundation Youth Fund Project, grant number (LQN25C130001).

---

**Fig. S1.**
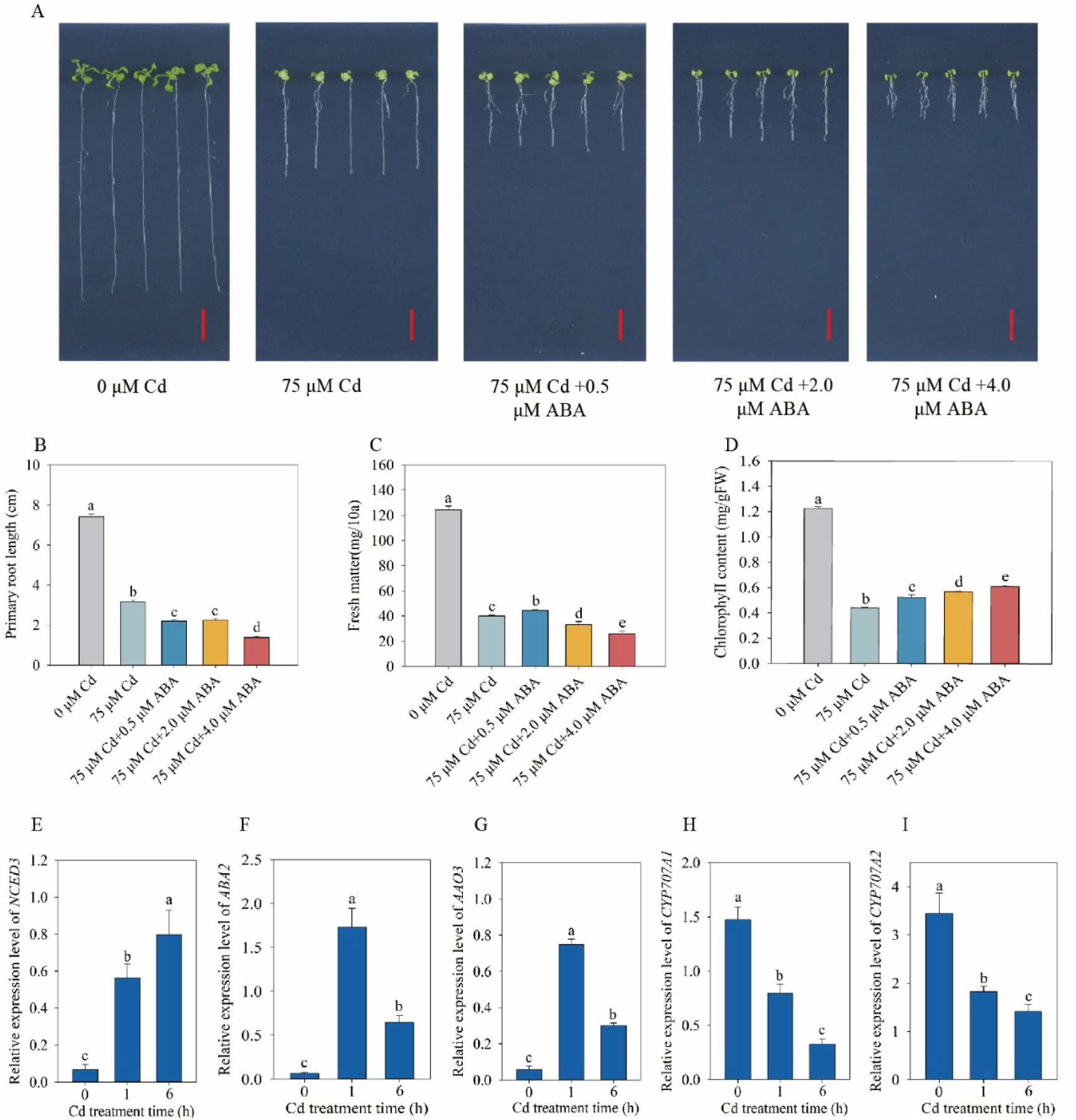
(A-D) Application of ABA enhances Cd tolerance of Arabidopsis. (A) Arabidopsis seedlings were grown on 1/2 MS media for 2 days, and then treated with 0 μM Cd, 75 μM Cd, 75 μM Cd+0.5 μM ABA, 75 μM Cd +2 μM ABA,75 μM Cd +4 μM ABA for 10 days, respectively. Scale bar, 1 cm. The primary root length (B), fresh weight (C) and chlorophyll content (D) of Col-0 were measured. Three independent experiments were done with similar results. Four plants per genotype from one plate were measured for each repeat. Data present the means ± SE (n = 4). Different letters indicate significant difference at *P <* 0.05 level (one-way ANOVA with Turkey’s test). (E-J) The relative expression level of *NCED3, ABA2, AAO3, CYP707A1, CYP707A2, ABI5* in Arabidopsis. qRT-PCR analysis of transcript levels of *NCED3* (E), *ABA2* (F), *AAO3* (G), *CYP707A1* (H), *CYP707A2* (I) and *ABI5* (J) in Arabidopsis respectively, at different time point after Cd stress. The relative expression levels of *NCED3, ABA2, AAO3, CYP707A1, CYP707A2* and *ABI5* was normalized against *ACTIN2* (*AT3G18780*). Data represent means ± SD (n = 4) from four independent experiments. Different letters indicate significant difference at *P <*0.05 (one-way ANOVA with Turkey’s test).

**Fig. S2.**
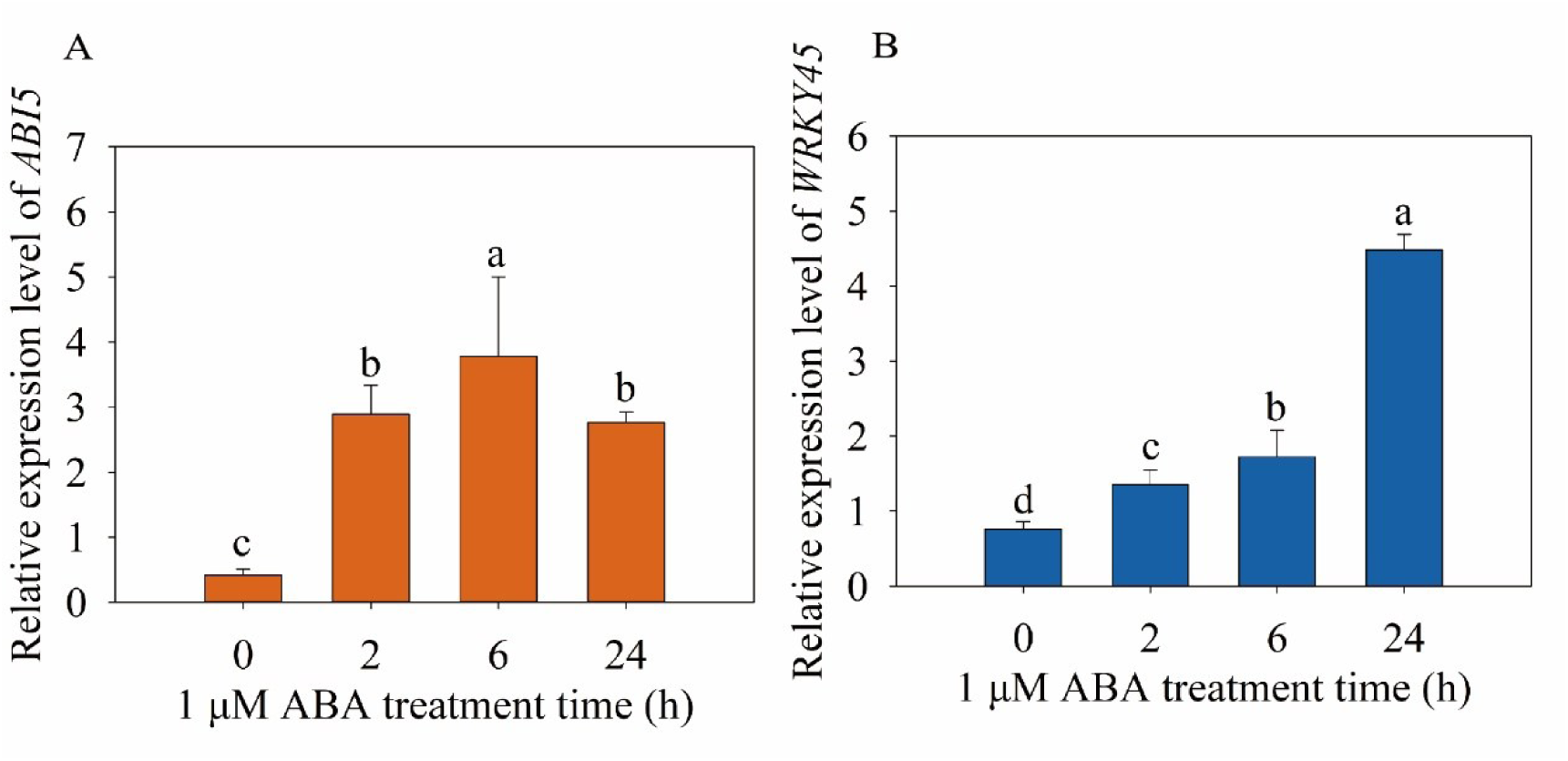
(A) The relative expression level of *ABI5* in Arabidopsis after 0, 2, 6, 24 h ABA treatment. The relative expression level of *ABI5* was normalized with *ACTIN2* (*AT3G18780*). Data represent means ± SD (n = 4) from four independent experiments. Different letters indicate significant difference at *P <*0.05 (one-way ANOVA with Turkey’s test).

**Fig. S3.**
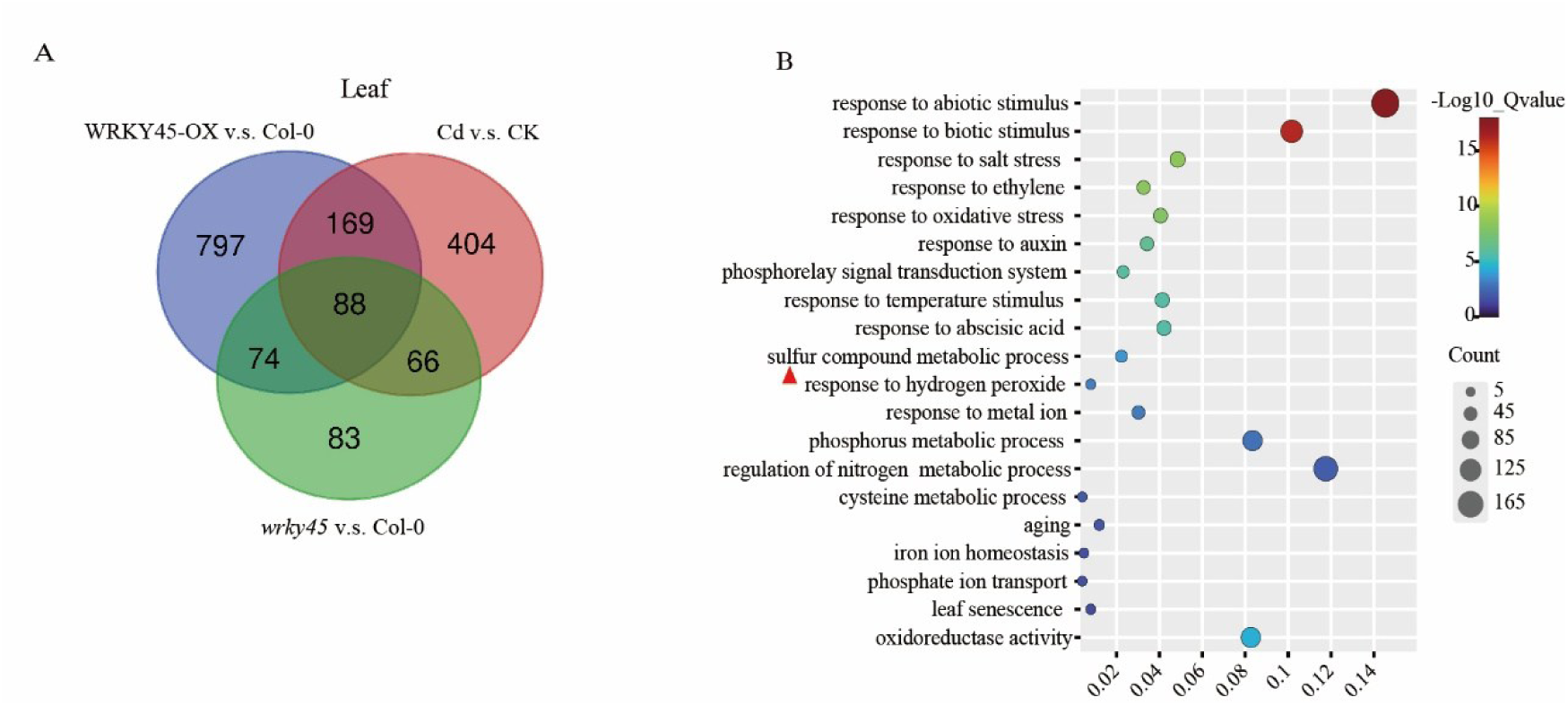
Venn and GO analysis of differential genes in leaf transcriptome sequencing of Col-0, *wrky45* and WRKY45OX-15 strains(A) Venn diagram illustrating the differentially expressed genes (DEGs) shared between Col-0, *wrky45* and WRKY45-OX-15 roots after 6 hours of Cd exposure. (B) GO term enrichment analysis of DEGs in the biological process in Col-0, *wrky45* and WRKY45-OX-15 roots after 6 hours of Cd exposure.

**Fig. S4.**
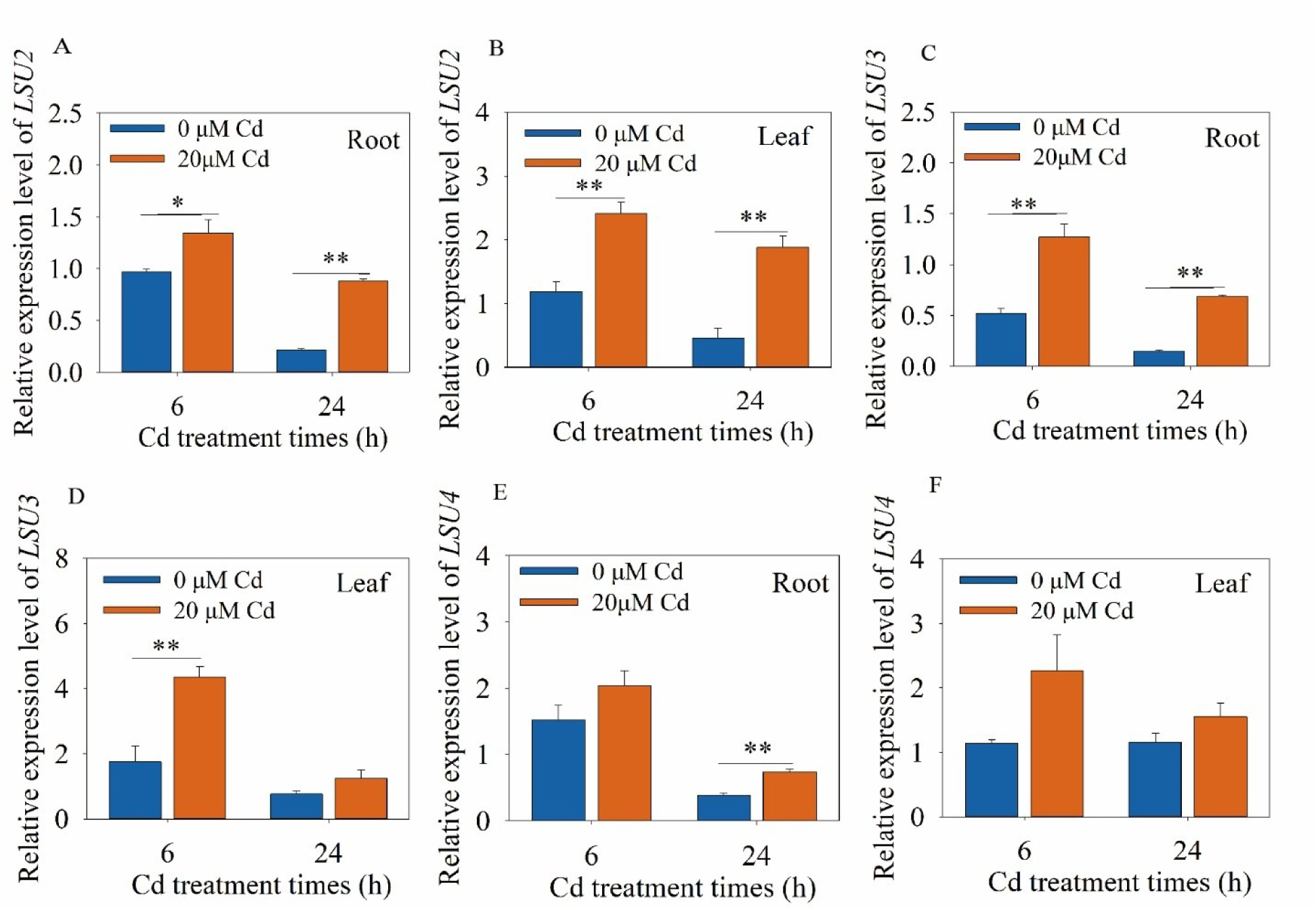
The relative expression levels of *LSU2* (A, B), *LSU3* (C, D) and *LSU4* (E, F) in leaves and roots of Arabidopsis. Hydroponic cultured three-week-old Arabidopsis seedlings were treated with 20 µM CdCl_2_ solution for 6 h and 24 h. The relative expression was normalized against *ACTIN2* (*AT3G18780*). Data are means ± SE (n = 4). The asterisk indicates that there is a significant difference between the control and the control (*, *P <* 0.05, **, *P <* 0.01; Student’s *t*-test).

**Fig. S5.**
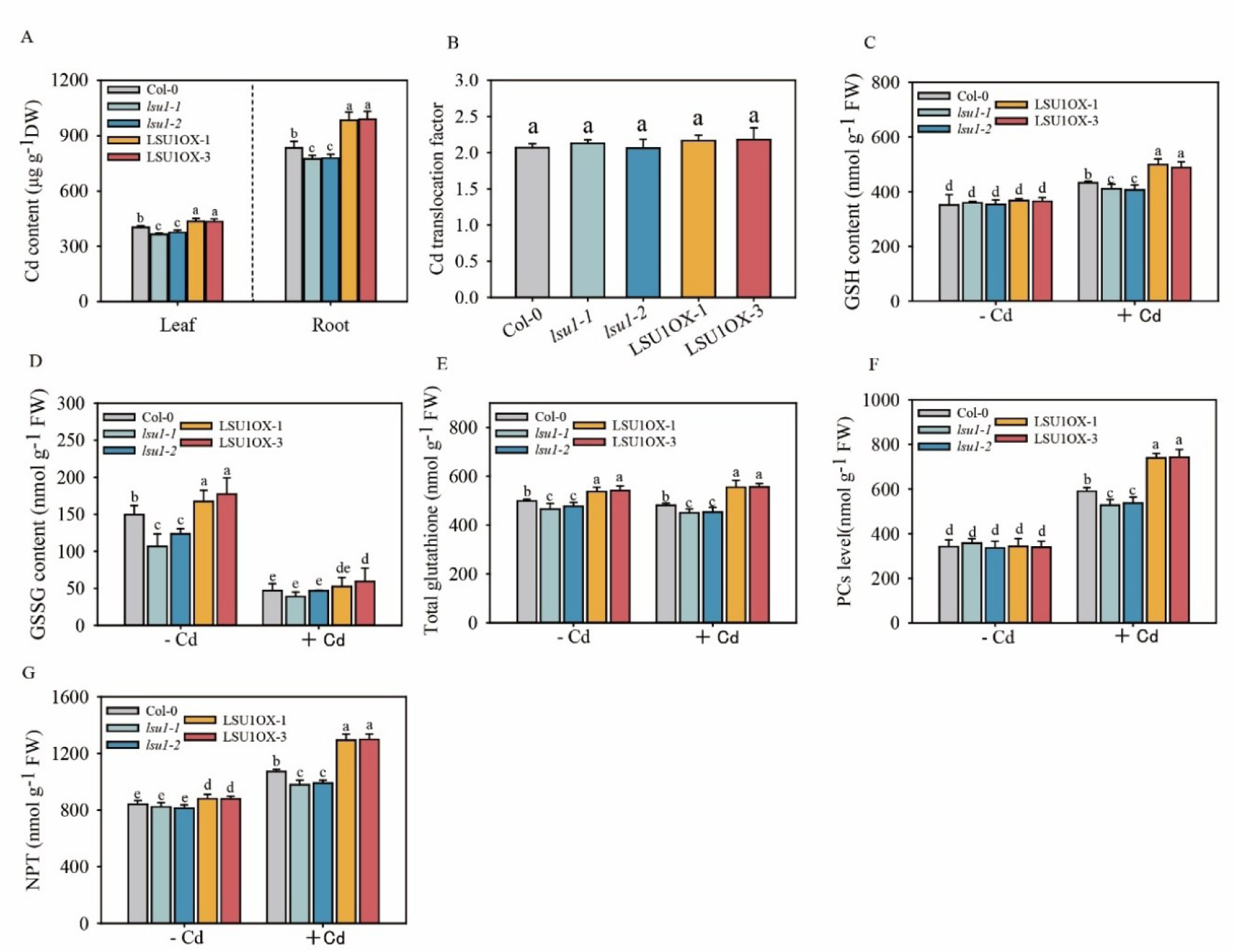
*LSU1* promotes the production of NBT, GSH and PC. Measurement of non-protein mercaptan (NPT) (A), total glutathione (B), PC (C), contents of Cd (D) and Cd translocation factor (E) in Col-0, *lsu1-1, lsu1-2*, LSU1OX-1 and LSU1OX-3 plants. The seedlings were cultured in nutrient solution for 2 weeks. After treated with 0 and 20 μM cadmium chloride for 24 h, the contents of NPT (A), GSH (B), PC (C) contents of Cd (D) and Cd translocation factor (E) were determined. Data present the means ± SE (n = 4). Different letters indicate significant difference at *P <* 0.05 level (one-way ANOVA with Turkey’s test).

**Fig. S6.**
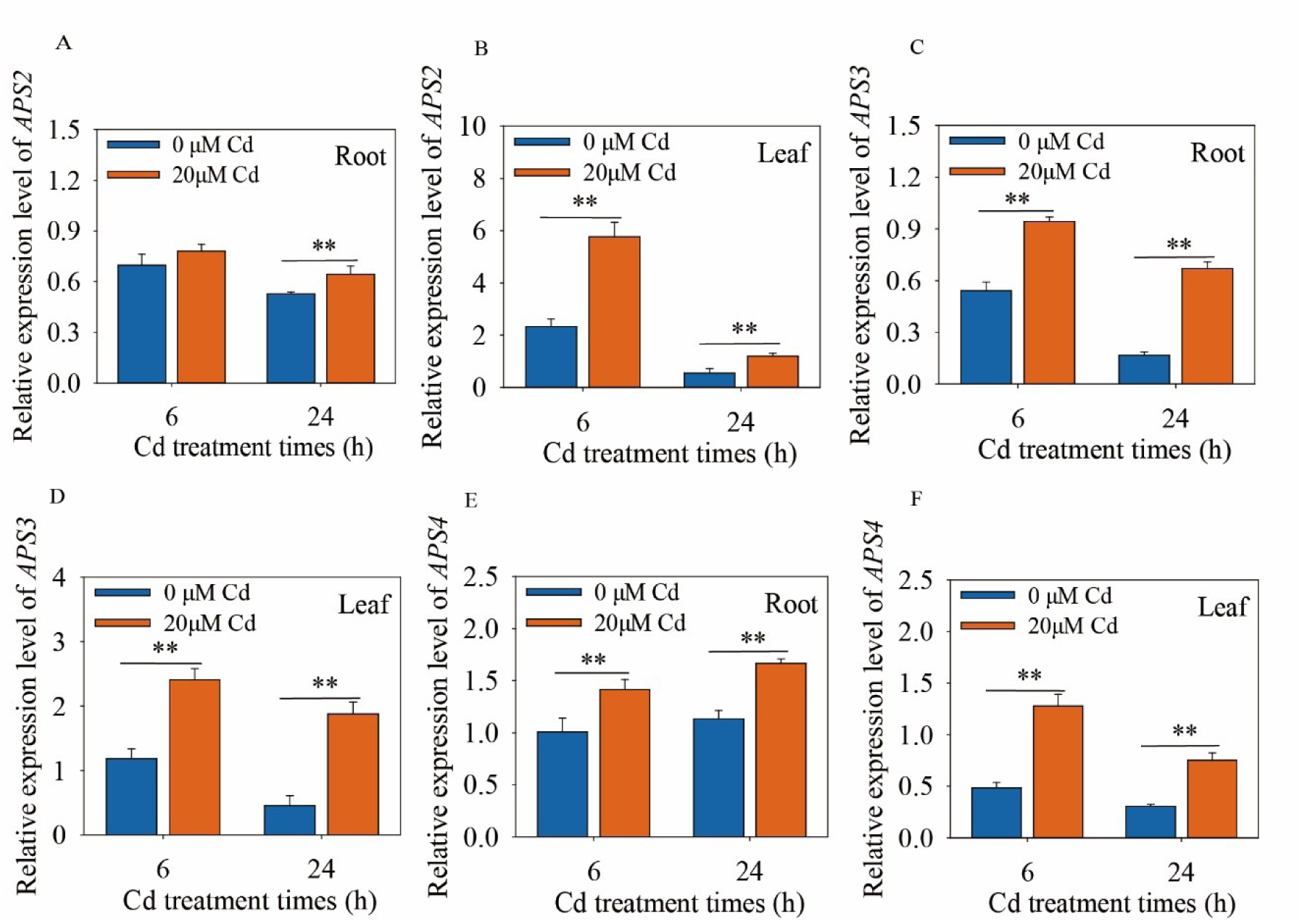
The relative expression levels of *APS2* (A, B), *APS3* (C, D) and *APS4* (E, F) in leaves and roots of Arabidopsis. Hydroponic cultured three-week-old Arabidopsis seedlings were treated with 20 µM CdCl_2_ solution for 6 h and 24 h. The relative expression was normalized against *ACTIN2* (*AT3G18780*). Data are means ± SE (n = 4). The asterisk indicates the significant difference between the control and treatment (*, *P <* 0.05, **, *P <* 0.01; Student’s *t*-test).

**Fig. S7.**
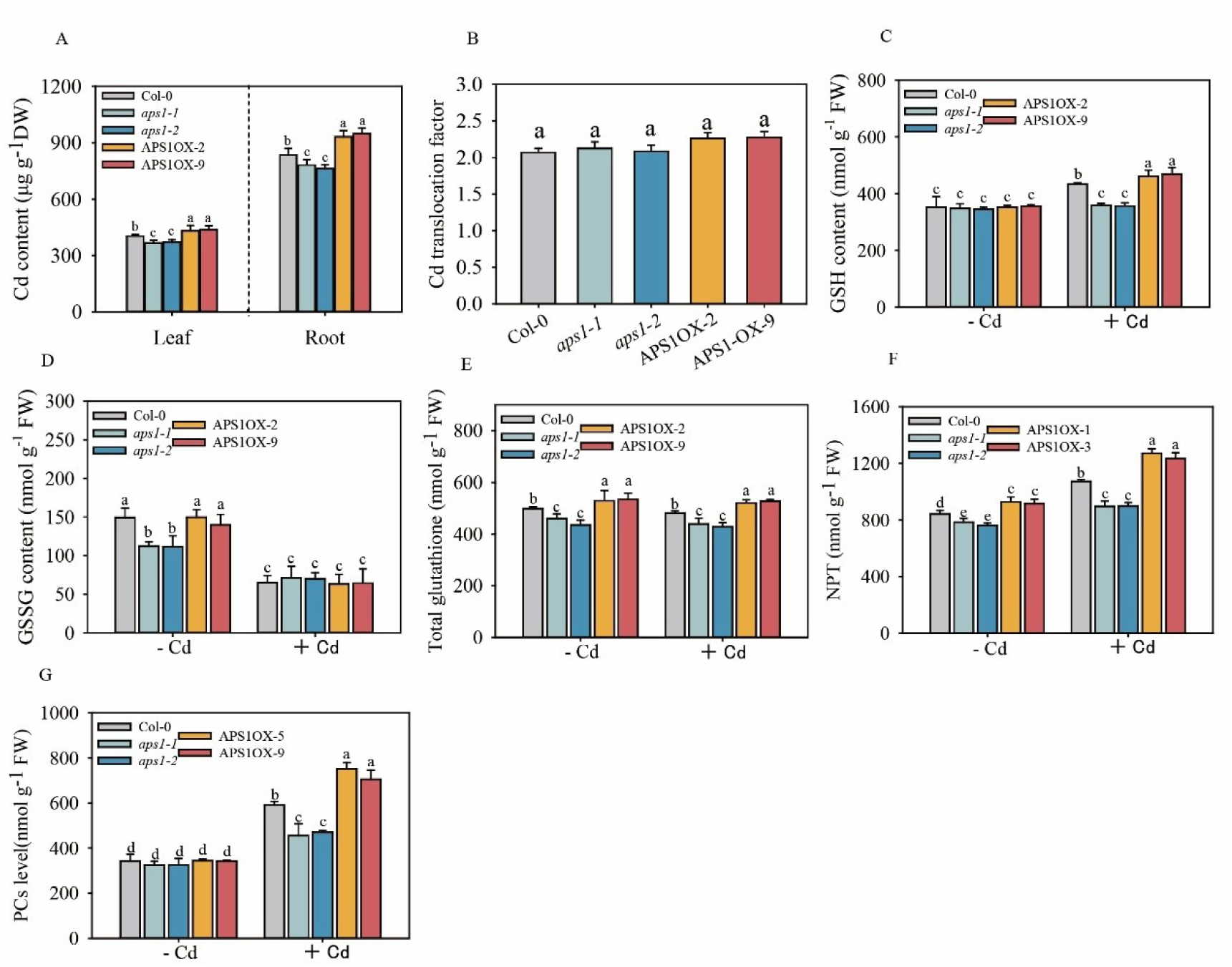
APS1 promotes the production of NBT, GSH and PC. Measurement of non-protein mercaptan (NPT) (A), total glutathione (B), PC (C), contents of Cd (D) and Cd translocation factor (E) in Col-0, *aps1-1, aps1-2*, APS1OX-2 and APS1OX-9 plants. The seedlings were cultured in nutrient solution for 2 weeks. After treated with 0 and 20 μ M cadmium chloride for 24 h, the contents of NPT (A), GSH (B), PC (C) and Cd (D), as well as Cd translocation factor (E) were determined. Data present the means ± SE (n = 4). Different letters indicate significant difference at *P <* 0.05 level (one-way ANOVA with Turkey’s test).

**Fig. S8.**
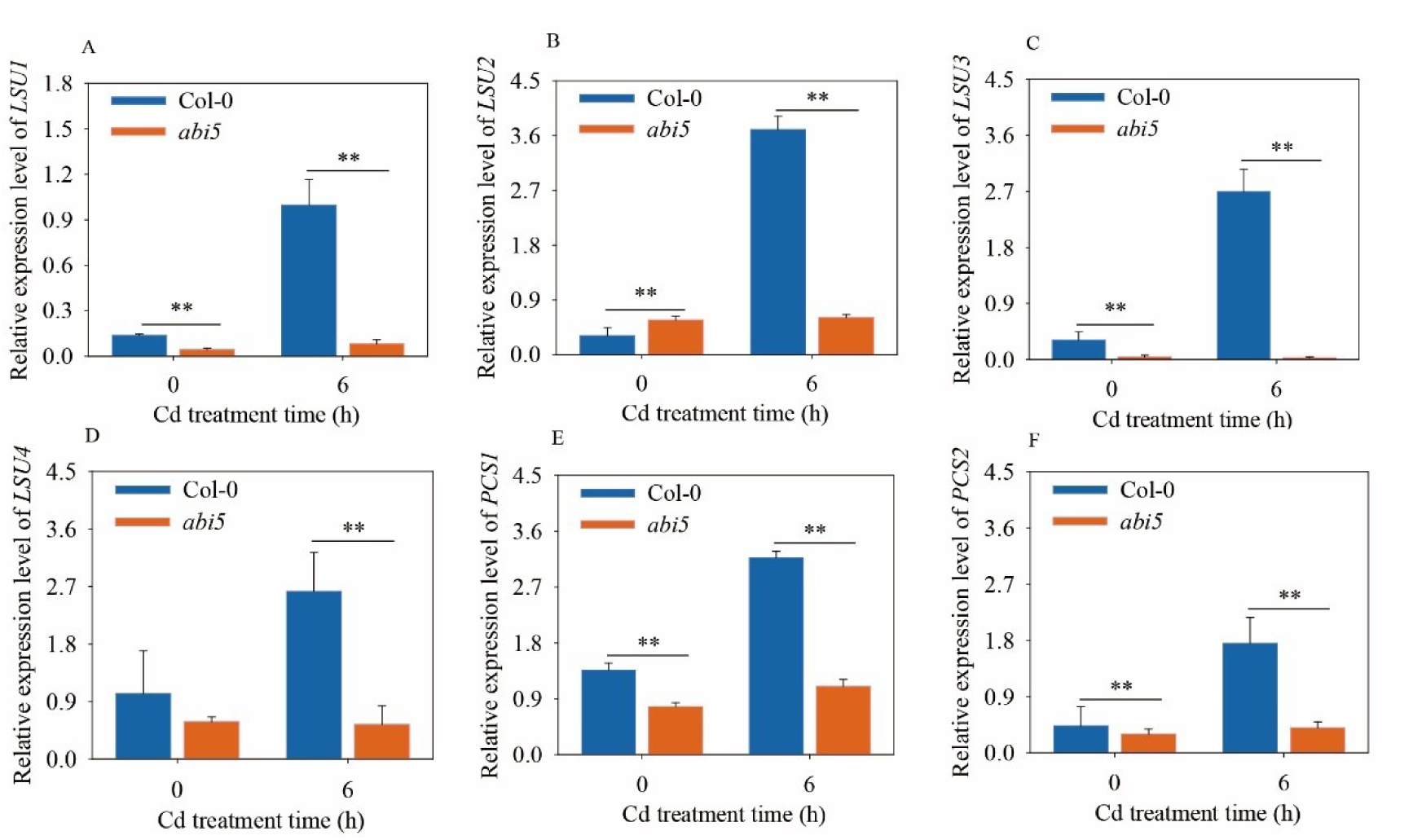
The relative expression level of *LSU1, LSU2, LSU3, LSU4 PCS1* and *PCS2* in Arabidopsis. qRT-PCR analysis showed the transcript levels of *LSU1* (A), *LSU2* (B), *LSU3* (C), *LSU4* (D), *PCS1* (E) and *PCS2* (F) in the wild type (Col-0) and *abi5-8* mutant respectively, at different time point after Cd stress. The relative expression levels of these genes were normalized against *ACTIN2* (*AT3G18780*). Data represent means ± SD (n = 4) from four independent experiments. The asterisks indicate significant differences from the control (*, *P <* 0.05, **, *P <* 0.01; Student’s *t*-test).

